# Synaptic synergy of T cell receptor and interleukin 2 receptor in CD8^+^ T cells

**DOI:** 10.1101/2024.08.13.607831

**Authors:** Jesusa Capera, Ashwin K Jainarayanan, Salvatore Valvo, Lina Chen, Steven N Quayle, Raymond J Moniz, Anish Suri, Michael L Dustin

**Affiliations:** The Kennedy Institute of Rheumatology, Nuffield Department of Orthopaedics, Rheumatology and Musculoskeletal Sciences, University of Oxford, Oxford, United Kingdom; Cue Biopharma, Boston, MA, USA

## Abstract

Interleukin 2 (IL2) promotes T cell proliferation and differentiation, making it a central target in immunotherapies. T cells fine-tune their sensitivity to and consumption of IL2 by regulating surface expression and composition of the IL2 receptor. Following antigen recognition, IL2 receptor signaling is shared through polarized interactions in T cell aggregates. However, how IL2 function is integrated during earlier antigen-dependent T cell synapses is unknown. Here, we demonstrate a synergistic effect between the T cell receptor (TCR) and IL2 receptor signaling at the immunological synapse of CD8^+^ T cells with supported lipid bilayers. TCR and IL2 signaling overlapped in space and time, potentiating each other when simultaneously triggered. Immuno-STATs, a safe and effective new class of immunotherapeutics, which fuse IL2 and peptide-major histocompatibility complex (pMHC) in a single molecule to expand antigen-specific CD8^+^ T cells, enhanced both TCR and IL2 signaling and promoted antigen specific T-T immunological synapses.

## INTRODUCTION

Interleukin 2 (IL2) is a 15 kDa cytokine primarily produced by activated CD4^+^ and, to a lesser extent, by activated CD8^+^ T cells. It acts as a potent autocrine/paracrine mitogenic signal and regulates the expansion and differentiation of the activated T cell subsets, crucial for immunity against cancers and infectious diseases (Spolski et al., 2018). IL2 also plays a critical role in the development and maintenance of regulatory T cells (Tregs) (Chinen et al., 2016). High-dose IL2 is an approved therapy for metastatic melanoma and renal cell carcinoma, with clinical benefit to a fraction of patients (Spolski et al., 2018). Despite its efficacy, the broad utility of IL2 for immunotherapy of cancers has been impeded due to toxicities related to indiscriminate activation of immune cells (Dutcher et al., 2014). To overcome these challenges, strategies aim to limit IL2 activity to tumour-specific immune cells, minimizing systemic effects and creating a "therapeutic index" for IL2—a clear separation between the lethal and effective doses. Approaches include generating IL2 variants with different receptor affinities (Levin et al., 2012), protease-sensitive masked IL2 molecules for local tumour-specific effects (Puskas et al., 2011), and more recently directing IL2 to T cells (Raeber et al., 2022, Moynihan et al., 2024). However, none of these approaches *selectively* target IL2 to tumour-specific T cells, which are the desirable cell population for therapeutic benefit and present in extremely low precursors frequencies within the polyclonal T cell repertoire (Ott et al., 2017). In this work, we study a new type of immunotherapeutic strategically designed to surmount this challenge. Immuno-STATs (IST) deliver IL2 to antigen-specific CD8^+^ T cells by combining affinity-reduced IL2 with pMHC molecules. This strategy goes beyond *cis*-targeting the IL2 receptor and the TCR, and directly exploits the natural synergy between the two signaling pathways to both create and functionalise homotypic T cell – T cell (T-T) immunological synapses and promote full activation and expansion of antigen-specific CD8^+^ T cells.

The relationship between IL2 and T-Cell Receptor (TCR) from the perspective of immunological synapse (IS) formation and signaling remains to be fully elucidated (Sabatos et al., 2008).The IL2 receptor consists of three transmembrane subunits, which bind to IL2 with varying affinities. The IL2RA (CD25) forms the low-affinity receptor. The IL2RB (CD122) and IL2RG (CD132) form the intermediate-affinity receptor, and all three form the high-affinity receptor. Only the IL2RB and IL2RG contain signaling cytoplasmic tails, primarily signaling via the Janus Kinase/Signal transducer and activator of transcription (JAK/STAT) pathway (Spolski et al., 2018, Ross et al., 2016, Malek, 2008, Kolios et al., 2021). IL2 binding to IL2RB/IL2RG recruits Jak1 and Jak3, leading to STAT activation and nuclear translocation to regulate gene expression (Villarino et al., 2011, Hu et al., 2021). The STAT family is composed of seven members and STAT5A and B are the more potently activated by IL2 receptors (Liao et al., 2013). IL2RG, also known as the common cytokine receptor γ chain (γc), is shared by IL4, IL7, IL9, IL15, and IL21 receptors, while IL2RB is shared by IL2 and IL15 receptors (Gonnord et al., 2018). Both CD4^+^ and CD8^+^ T cells transiently express high-affinity IL2 receptors following TCR engagement, peaking within hours post-antigen challenge and declining after 24 hours (Malek, 2008). The strength of IL2 receptor signaling influences CD8^+^ T cell differentiation, with high IL2RA expression driving effector cell proliferation, while lower IL2RA expression promotes memory cell formation (Manjunath et al., 2001, Spolski et al., 2018, Kalia et al., 2010). Additionally, IL2 shapes the effector CD8^+^ T cell repertoire, promoting stemness-like features through TCF-1 expression and preventing terminal exhaustion (Codarri Deak et al., 2022, Hashimoto et al., 2022).

IL2 directly affects CD8^+^ T cells sensitivity to antigenic peptides, with CD4^+^ effector T cells potentially releasing IL2 to CD8^+^ T cells as they synapse with dendritic cells (DC) (Au-Yeung et al., 2017). However, IL2 functions primarily through paracrine or autocrine signaling following T cell-DC interaction, with activated T cells forming large, dynamic clusters sustained by LFA-1 interactions (Gerard et al., 2013),(Zenke et al., 2020). These clusters exhibit directional IL2 secretion and polarization of the IL2 receptor complex towards T-T synapses, enhancing pSTAT5 signaling and IL2 action (Sabatos et al., 2008). The T-DC IS can be modelled using supported lipid bilayers (SLBs), which simplifies the IS into a 2D model with concentric compartments for effector function (central), adhesion and transport (peripheral) and signal initiation/costimulation (distal ring/corolla) (Demetriou et al., 2020). F-actin dependent TCR microclusters are sites of TCR signaling (Yokosuka et al., 2008, Bunnell et al., 2001) and are analogous to microvilli and podosomes in 3D models (Varma et al., 2006, Kumari et al., 2015, Cai et al., 2017). IL2 roles in the IS between CD4^+^ or CD8^+^ T cells and DCs, also involve Tregs, which intercept IL2 to enhance their survival while starving autoreactive T cells, impacting the fitness of these cells (Wong et al., 2021). This dual function of IL2 is critical for immunotherapy, where targeted delivery of IL2 must secure a balance between its immunosuppressive and pro-inflammatory effects to maximize therapeutic benefits in specific disease contexts.

In this report, we describe the co-localization of IL2 receptor subunits and signaling at the IS triggered by pMHC-specific TCR engagement at different stages of CD8^+^ T cell differentiation. The natural synergy between the TCR and IL2R signaling forms the mechanistic basis for a new class of T cell engagers, ISTs, which direct IL2 activity towards tumour-specific CD8^+^ T cells to selectively expand them (Quayle et al., 2020, Seidel et al., 2021). This selective activation creates a therapeutic index for IL2, i.e. favouring biased activation of tumour-specific T cells while avoiding the systemic activation of other immune cells. To note, lead IST clinical candidates, CUE-101 and CUE-102, are currently being evaluated in clinical trials of solid cancers, showing efficacy via enhancing survival in late-stage metastatic cancer patients and significantly increasing response rates in combination with anti-PD-1 mAb in frontline metastatic patients (Zhang et al., 2021). The current ongoing clinical trials with ISTs have provided evidence of clinical proof of concept and de-risking via demonstration of tolerability (i.e., no major IL2-related toxicities) and anti-tumour efficacy (Zhang et al., 2021).

## RESULTS

### Heterogeneous expression and distribution of the IL2 receptor subunits at the IS

T cell responsiveness to IL2 relies on the expression of IL2 receptor subunits and other interleukin receptors competing for the common-gamma chain (Spolski et al., 2018, Zhou et al., 2022). Therefore, the distribution of the IL2 receptor subunits was analysed at the IS of three different states of CD8^+^ T cell differentiation: naïve, memory and activated lymphoblast (blast). First, expression of the IL2 receptor subunits was confirmed using a publicly available sc-RNAseq dataset. UMAP analysis established a distinct cluster of CD8^+^ T cells with greater heterogeneity in memory cells and blasts (Fig. 1A). IL2RA was primarily detected in blasts (Fig. 1B, C). IL2RB was well expressed in memory and blast CD8^+^ T cells and nearly absent in the naive state, while IL2RG was well expressed across all states (Fig. 1B, C). Expression of other interleukin receptor subunits competing for the γ_c_ (IL2RG) was also analysed. The IL7R was mainly expressed in the naïve and memory states, with markedly decreased expression in blasts (Fig. 1C).

**Figure 1:**
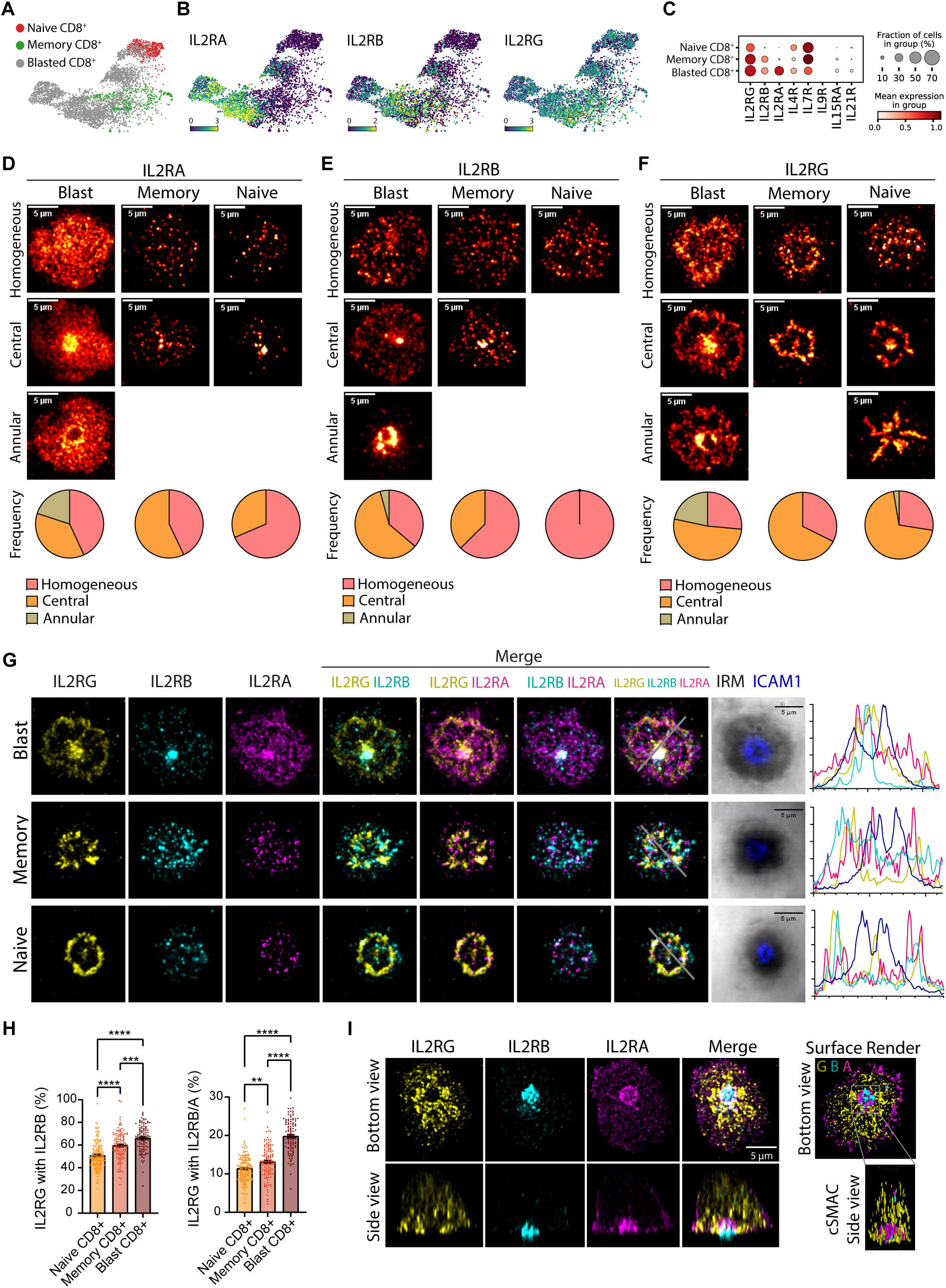
Distribution of the IL2 receptor complex at the IS. **(A)** UMAP dimensionality reduction embedding of blasted (grey), memory (green) and naive (red) CD8^+^ T cells coloured by orthogonally generated clusters labelled by manual cell type annotation. **(B)** UMAP embedding of scRNAseq dataset of CD8^+^ T cell populations coloured by expression profiles of IL2RA (left), IL2RB (middle) and IL2RG (right). **(C)** Dot plots depicting mean expression (visualized by colour) and fraction of cells (visualized by the size of the dot) expressing common-gamma chain interleukin receptor genes. **(d-f)** RA14 CD8^+^ T cells were exposed to Ag-specific SLBs in the presence of 50 U/ml of IL2 to form ISs and fixed after 10 min. Samples were then permeabilized and immunostained against the IL2 receptor subunits. Panels show representative TIRF images of the different distribution phenotypes (Homogeneous, Central and Annular) observed for the IL2RA (d), IL2RB (e) and IL2RG (f) for each state of CD8^+^ T cells (Blast, Memory and Naïve). Frequency for each phenotype is represented as parts of a whole for each T cell state. Data is the mean of three independent donors with >50 cells/donor. **(G)** Representative TIRF images of blast (top), memory (middle) and naïve (bottom) RA14 CD8^+^ T cells forming ISs with Ag- specific SLBs in the presence of 50 U/ml of IL2 and fixed after 10 min. Samples were then permeabilized and immunostained for the IL2 receptor subunits. Histograms (right) show the intensity line profile for the different proteins. **(H)** Percentage of the total IL2RG intensity that is within the area covered by IL2RB (left plot) or within the intersection between the IL2RB and IL2RA (right plot). See Fig. S3 for a more detailed explanation. Dots shows the distribution of the values for >100 cells, from three independent donors. Histograms show the mean +/- SE. **pv<0.01, ***pv<0.001, ****pv<0.0001; Kruskal-Wallis test. **(I)** Representative z-stack Airyscan reconstruction of a RA14 CD8^+^ blast forming a IS on an Ag-specific SLB in the presence of 50 U/ml of IL2. Sample was fixed after 10 min of SLB exposure, permeabilized and immunostained against the IL2 receptor subunits.

Next, the distribution of the IL2 receptor subunits was visualized at the IS of antigen (Ag)-specific CD8^+^ T cells. CD8^+^ T cells were electroporated with mRNA encoding the CMV pp65 RA14 TCR and exposed to supported lipid bilayers (SLBs) presenting HLA-A201 complexes with the CMV pp65_495- 502_, in addition to ICAM1, CD80 and CD58. The T cells were given 10 minutes to interact with the SLBs at 37°C prior to fixation and imaging with total internal reflection fluorescence microscopy (TIRFM), which isolates fluorescence signals within 200 nm of the SLB. IL2 receptor subunits displayed punctate elements in an overall pattern that could be classified into *homogeneous* (puncta evenly distributed throughout the synaptic contact), *centralized* (dense clustering of puncta at centre) and *annular* (a central ring of puncta) (Fig. 1D-F, Fig. S1).

IL2RA displayed the strongest signal in blasts, as expected, but was also detected in memory and naïve T cells (Fig. 1D, Fig. S2A). IL2RA was centralized in most of the blast and memory ISs, while the few puncta visible in naïve T cells appeared homogenously scattered in the IS (Fig. 1D). IL2RB was mostly centralized in blasts, but homogenous patterns were more prominent in memory cells and dominant in naïve cells (Fig. 1E). IL2RG was mainly centralized in all T cell states and formed a peripheral corolla, which was observed as a secondary feature in all types of CD8^+^ T cell states (Fig. 1F).

The centralized phenotypes of IL2 receptor subunits were more frequent when the IS was established in the presence of IL2 (Fig. S1). Simultaneous visualization of the three subunits at the IS showed distinct localizations, with some puncta containing two or all three subunits. (Fig. 1G). The abundance of the subunits within these clusters was dependent on the T cell differentiation state. For example, blast cells showed the highest percentage of IL2RG within IL2RB or IL2RB/A clusters (potential intermediate or high affinity IL2 receptors, respectively), followed by memory cells (Fig. 1H and Fig. S3). This pattern is inverse to the expression of other common-gamma chain receptors like IL7R (Fig. 1C). Pairwise correlations were highest in blasts and lower for memory and naïve CD8^+^ T cells (Fig. S3E).

Three-dimensional imaging of cells forming IS with the SLBs indicated that centrally accumulated subunits were partly above the plane of the SLB, suggesting internalization or release through ectocytosis of extracellular vesicles or secretion in exosomes (Kvalvaag et al., 2023). IL2RA and IL2RB were partly segregated, while IL2RG was abundant in intracellular compartments away from the IS (Fig. 1I).

### IL2/IL2 receptor subunits are released through extracellular vesicles

To further investigate whether IL2 receptor subunits were released in extracellular vesicles (EVs), mass spectrometry (MS) data from EVs released by human CD8^+^ T cells at different differentiation states (naïve, memory or total/pooled CD8^+^ T cells) were analysed. EVs were purified before (resting) and after (activated) treatment with anti-CD3/CD28 Dynabeads (Jainarayanan et al., 2023). In resting CD8^+^ T cells, only IL2RG was present in EVs, along with EV marker CD81, CD8 and CD3E (Fig. 2A). In contrast, EVs from activated CD8^+^ T cells contained IL2, IL2RA and IL2RB, in addition to detectable levels of IL2RG, IL7R and IL9R (Fig. 2A). Data was validated by Nano-flow cytometry, which showed that resting EVs were 7.5-times more likely to be IL2RG single positive than activated EVs, whereas activated EVs were 10-times more likely to be single positive for IL2RB than resting EVs. Double positive EV were rare, suggesting that EV formation operates on the individual receptor subunits rather than on intact complexes (Fig. 2B-C).

**Figure 2:**
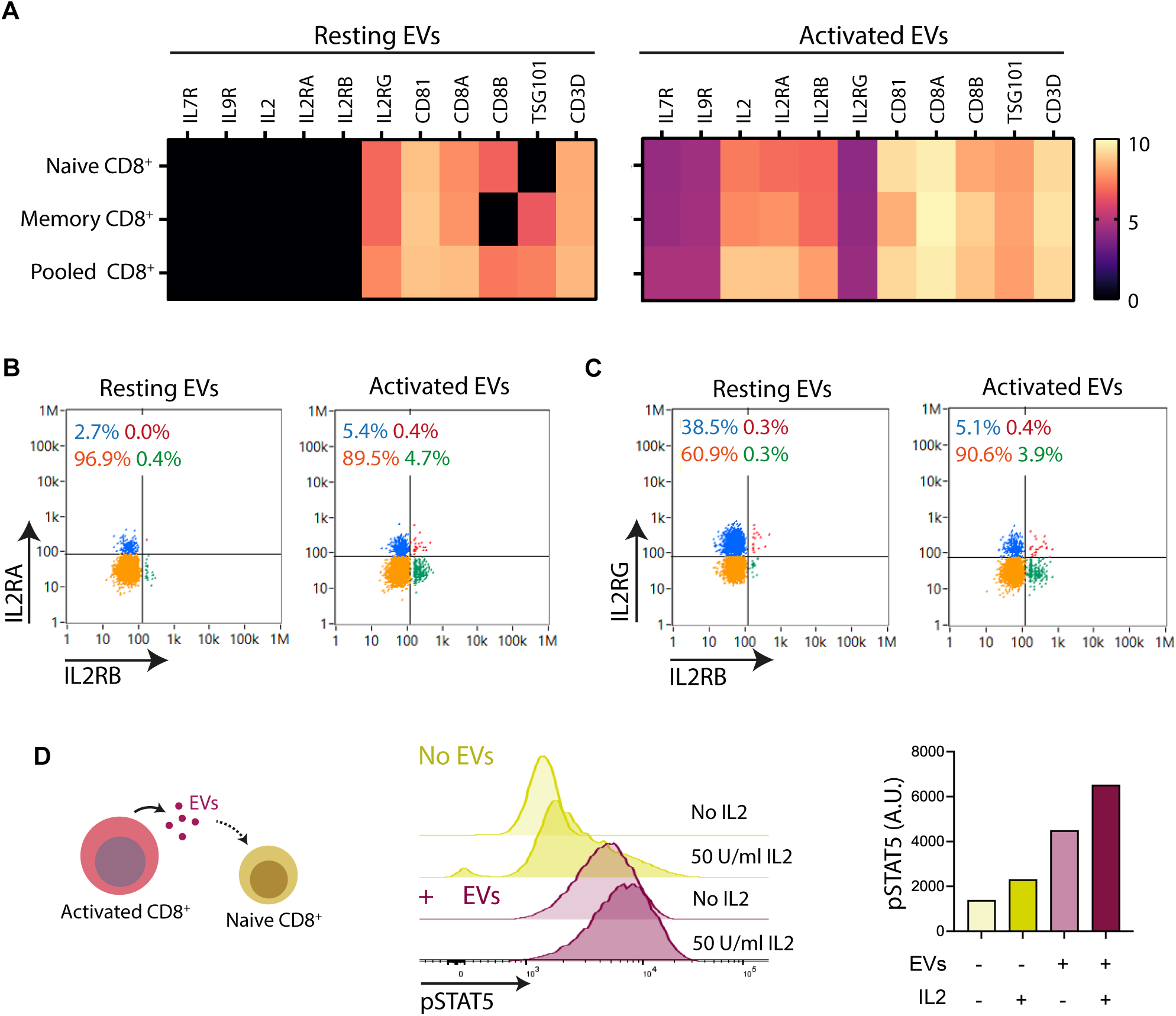
Release of the IL2 receptor subunits in EVs is dependent on the activation state of the CD8^+^ T cells. **(A)** Heatmap showing the normalized expression of proteins identified in EVs from resting and activated CD8^+^ T cells (naïve, memory and total/pooled CD8^+^ differentiation states). All members of the common-gamma chain cytokine receptors and interleukins were interrogated, together with other activation markers and EVs markers. The heatmap shows all species detected. For activation, cells were incubated with anti-CD3/CD28 Dynabeads for 48 hours. **(B)** Nano-FCM analysis showing the scatter plot for IL2RA (y-axis) and IL2RB (x-axis) positive EVs from resting and activated pooled CD8^+^ T cell. Marked in red are the double positives (IL2RA and IL2RB), blue exclusively represents the IL2RA positive EVs (negative for IL2RB), green exclusively represents IL2RB positive EVs (negative for IL2RA), and the double negatives are marked in yellow. Similarly, **(C)** shows the scatter plot for IL2RG (y-axis) and IL2RB (x-axis) positive EVs from resting and activated CD8^+^ T cell. Marked in red are the double positives (IL2RG and IL2RB), blue exclusively represents the IL2RG positive EVs (negative for IL2RB), green exclusively represents IL2RB positive EVs (negative for IL2RA), and the double negatives are marked in yellow. **(D)** Naïve CD8^+^ T cells were exposed to EVs purified from activated CD8^+^ T cells. After 48 hours with or without EVs incubation, Naïve CD8^+^ T cells were treated or not with 50 U/ml of IL2 and analysed by phospho-flow cytometry to quantify total pSTAT5 levels. Right plot shows the median intensity for pSTAT5.

Next, we evaluated if EVs from activated CD8^+^ T cells, which collectively contain IL2 and all the IL2 receptor subunits, can activate pSTAT5 in naive CD8^+^ T cells, which have low sensitivity to IL2. Note that size exclusion chromatography used to isolate the EV removes free IL2 (Jainarayanan et al., 2023). There was a robust increase in phosphorylation (pSTAT5) by naive CD8^+^ T cells treated with the activated EVs, which was greater than what could be achieved with soluble 50 U/ml of IL2 (Fig. 2D). Thus, IL2 and IL2 receptor components are released in EV that can trigger STAT5 activation in otherwise IL2 unresponsive naïve CD8^+^ T cells.

### Distribution of IL2 receptor activity at the IS

Responsiveness to IL2 is dependent on the CD8^+^ T cell state of differentiation (Fig. 1) (Gonnord et al., 2018). Upon IL2 treatment, naïve CD8^+^ T cells showed limited pSTAT5 compared to the memory or blasted counterparts, the later showing the highest pSTAT5 levels (Fig. 3A). In Ag-specific ISs formed by CMV-specific CD8^+^ T cell blasts on SLBs, pSTAT5 signaling was polarized towards the IS, even without soluble IL2 in the media (Fig. 3B). Puncta of pSTAT5 and pZAP70 appeared immediately upon synaptic contact, peaking within 2-15 min and returning near basal levels after 30 min (Fig. 3C-F). In addition, these signaling events initially overlapped at the IS but dissipated at later time points (Fig. 3C- F). pSTAT5 puncta, like pZAP70, also overlapped with TCR microclusters (Fig. 3G-K).

**Figure 3:**
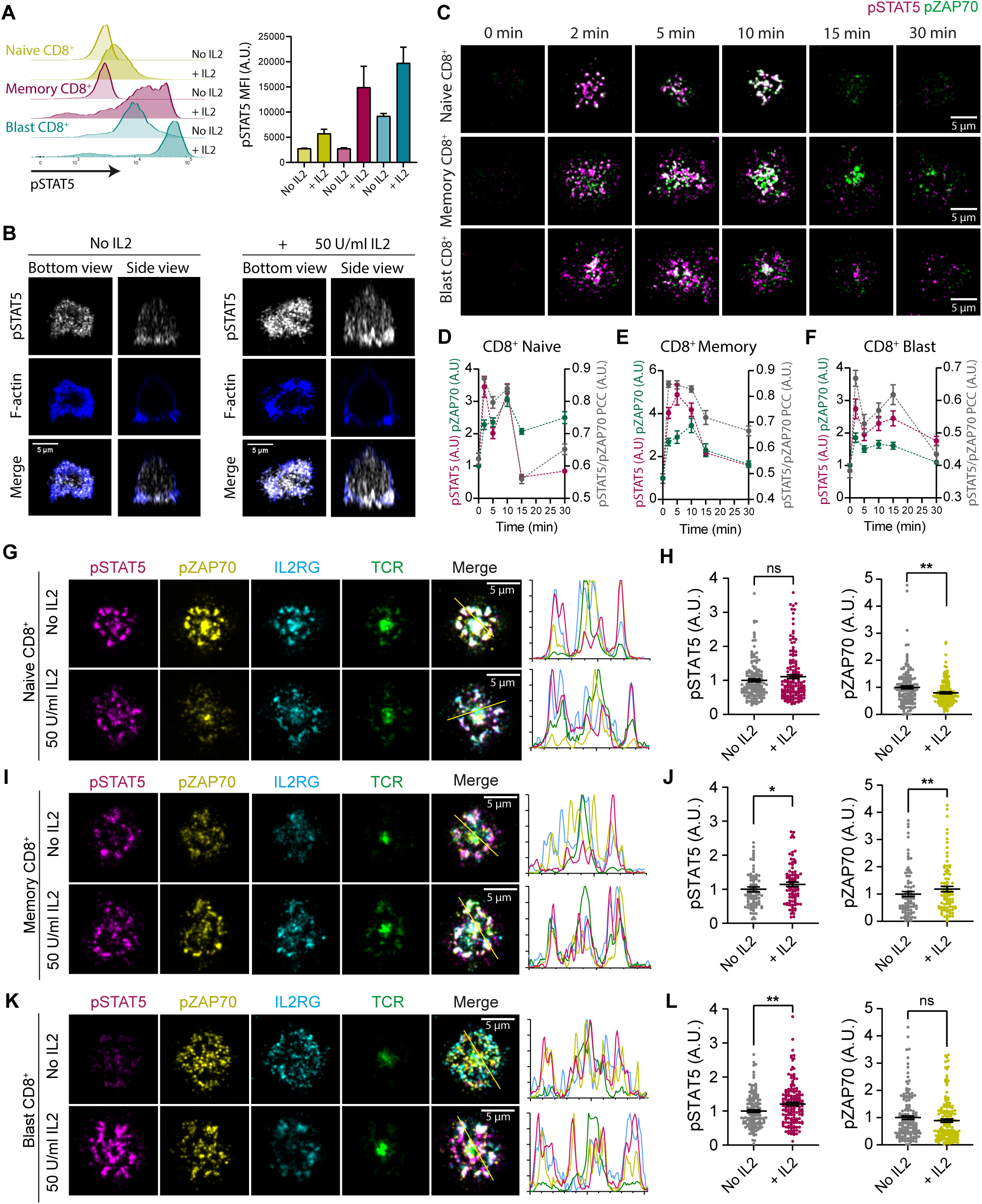
Distribution of IL2 receptor activity at the IS. **(A)** Histograms of phospho-flow cytometry data showing total levels of pSTAT5 after 10 min incubation at 37 ° C in the presence (+IL2) or absence (No IL2) of 50 U/ml of IL2 treatment; for naïve (yellow), memory (magenta) and blast (blue) CD8^+^ T cells. Bar plots (right) show the mean +/- SE of the pSTAT5 median fluorescence intensity (MFI) from three independent donors. **(B)** Representative z-stack Airyscan reconstruction of RA14 CD8^+^ blasts forming a IS on an HLA-A2-p65 peptide bearing SLB in the presence or absence of 50 U/ml of IL2. Sample was fixed after 10 min of SLB exposure, permeabilized and stained for F-actin and immunostained against pSTAT5. **(C)** Representative TIRF images of Naïve (top), Memory (middle) and Blast (bottom) CD8^+^ T cells forming an IS with a SLB and fixed after the indicated time-point. 0 min represents cells fixed after 15 min of contact with a SLB without UCHT1 (no IS formation). pSTAT5, magenta; pZAP70, green. White pixels show overlap between pSTAT5 and pZAP70. **(D-F)** Plots show the mean intensity for pSTAT5 (magenta) and pZAP70 (green) normalized to the 0 min time-point (left Y axis) and the Pearson’s Correlation Coefficient (PCC, grey) between pSTAT5 and pZAP70 signals (right Y axis) over time for naïve (d), memory (e) and blast (f) CD8^+^ T cells. Data is the mean +/- SE of 25-50 cells. **(G-K)** Naïve (g-h), memory (i-j) or blast (k-l) RA14 CD8^+^ T cells were exposed to Ag-specific SLBs in the presence or absence of 50 U/ml of IL2 to visualize IS formation. Cells were fixed after 10 min of bilayer exposure and immunostained for pSTAT5 (magenta), pZAP70 (yellow), IL2RG (cyan), TCR (green). (g, i and k) Representative TIRF images of ISs. Merge is the overlap between channels. Histograms on the right show the intensity profile across the yellow line shown in the merge images. (h, j and l) show the dot plots with the mean intensity for pSTAT5 and pZAP70 normalized to the condition without IL2. Data is the mean +/- SE, n=90-150 from 3 independent donors (50 cells/donor for naïve and blast and 30 cells/donor for memory). ns, not significant, *pv<0.05, **pv<0.01; Mann-Whitney test.

Adding IL2 in the media concomitantly to SLB exposure did not affect pSTAT5 levels in naïve CD8^+^ T cells (Fig. 3G,H). However, in both memory (Fig. 3I, J) and blast (Fig. 3K, L) cells IL2 boosted the pSTAT5 intensity. IL2 only increased the pZAP-70 signal in memory T cells (Figure 3I, J). The pSTAT5 and pZAP-70 co-localization in the IS was observed in the IS centre and the corolla (Figure 3G, I, K). Notably, the IS activated pSTAT5 through TCR and co-stimulatory signals, suggested by the presence of pSTAT5 punta even in the absence of exogenous IL2 or in the naïve CD8^+^ T cells, which respond weakly to IL2 (Fig. 3).

To further investigate the effects of IL2 on synapse formation, recruitment of TCR, ICAM1/LFA1, CD58/CD2 and CD80/CD28 to the synapse was evaluated in naïve, memory and blast CD8^+^ T. IL2 did not show major effects synapse assembly and the formation of the SMAC compartments was normal. However, in blast CD8^+^ T cells, IL2 enhanced the recruitment of signaling proteins to the IS, especially the CD2/CD58 pair, in a dose-dependent manner (Fig. S4). The synapse contact area was also larger in the presence of IL2 for naïve and blast CD8^+^ T cells (Fig. S5).

### Immuno-STATs (IST) induce Ag-specific IS

To further understand the relationship between IL2 receptor and TCR and its therapeutic potential, we incorporated the novel ISTs therapeutics into our SLB system. ISTs consist of a human IgG1 Fc domain, with mutations attenuating Fc receptor binding, fused to two copies of a HLA-A2 complex, a covalently tethered peptide epitope and four human IL2 variant molecules (CueIL2), which have reduced affinity for IL2RA and IL2RB subunits (H16A and F42A mutations) (Fig. 4A). As a negative control for the effects of IL2, a variant of the ISTs without IL2 molecules was designed (noIL2). In addition, both IST constructs (with and without IL2) were produced with two different types of peptide epitopes bound to the HLA-A201: cytomegalovirus pp65_495-502_ peptide (CMV), which is recognized by the RA14 TCR expressed by the T cells used in this work, and the MART1_26-35_ peptide (Mart1), which was used as a negative control (Fig. 4A). A synthetic Staphylooccus aureus protein A derived peptide (D-peptide) with human IgG binding activity was used to tether the ISTs to the SLB though the Fc.

We tested the ability of ISTs to mediate adhesion of T cells to the SLB without any additional adhesion or co-stimulatory molecules. RA14 CD8^+^ T cell blasts adhered to SLB presenting a high density (∼2000/µm^2^) of the CMV IST with or without the CueIL2 (Fig. 4B-C). No adhesion was observed with Mart1 IST regardless of IL2 presence (Fig. 4B-C). Notably, cell contacts were larger in size when CueIL2 was presented by the IST (Fig. 4D). The SLB-anchored ISTs were recruited to the cell-SLB contact and clustered to the centre of the contact even without adhesion and costimulatory molecules (Fig. 4E).

When ICAM1 molecules were added to the SLB at a lower IST density, typical IS formation with adhesion rings was confirmed only CMV peptide-presenting ISTs, and not the Mart1 peptide IST (Fig. 4F). Thus, the peptide-HLA component of the IST controls adhesive avidity independently of the attenuated IL2, and IS formation occurred only when the IST was presenting a cognate peptide for the TCR.

### Co-delivery of IL2 and antigen enhances IL2 receptor assembly and TCR complex proximity

The recruitment and distribution of the IL2 receptor complexes and the TCR was visualized in RA14 CD8^+^ T cell blasts forming ISs with SLB containing the different IST constructs (Mart1-noIL2; CMV- noIL2; Mart1CueIL2; CMVCueIL2) with ICAM-1, CD80 and CD58. The greatest IL2 receptor subunit recruitment occurred with IST constructs containing CueIL2 (Fig. 5A).

**Figure 4.**
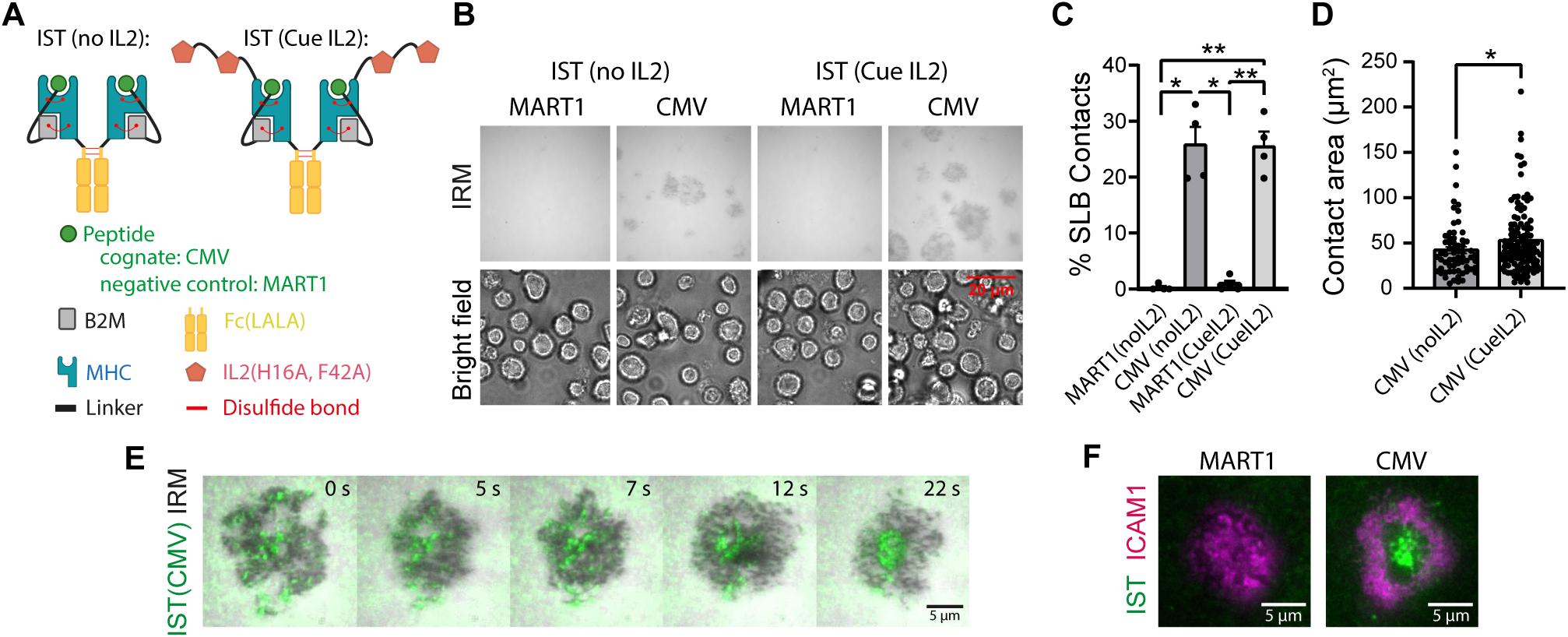
IST induced adhesion and IS formation. **(A)** Cartoon representing the different structural variants of IST used in this work. Two HLA-A*02 complexes with the cognate Ag (CMV pp65) or a non-specific control Ag (MART-1) are connected, or not, to four units of engineered IL2 (Cue IL2), with abrogated binding to IL2RA and reduced binding to IL2RB. **(B)** RA14 CD8^+^ blasts were exposed to SLBs containing the different structural variants of ISTs linked to the D-peptide at SLB-saturating conditions without any additional adhesion or co-stimulatory molecules and imaged by interference reflection microscopy (IRM) and bright field after 15 min. **(C)** Percentage of cells forming a contact with the SLB (detected by IRM) from all cells exposed to the SLB (bright field) in (b). Data is the mean +/- SE of 4 independent donors; *pv<0.05, **pv<0.01, one-way ANOVA test. **(D)** Quantification of the contact area between the SLB and the T cell (detected by IRM) from cells in (b). Data is the mean +/- SE of >60 established contacts. *pv<0.05; Mann-Whitney test. **(E)** Time-lapse of a RA14 CD8^+^ blast T cell forming a contact with a SLB containing fluorescently tagged ISTs with the CMV peptide (IST, green; IRM, grey). **(F)** RA14 CD8^+^ blast T cells were exposed to SLBs containing CMV-specific or MART1-specific ISTs (green) linked to the D-peptide at 3nM and ICAM1 (magenta) and imaged after 15 min of exposure.

**Figure 5.**
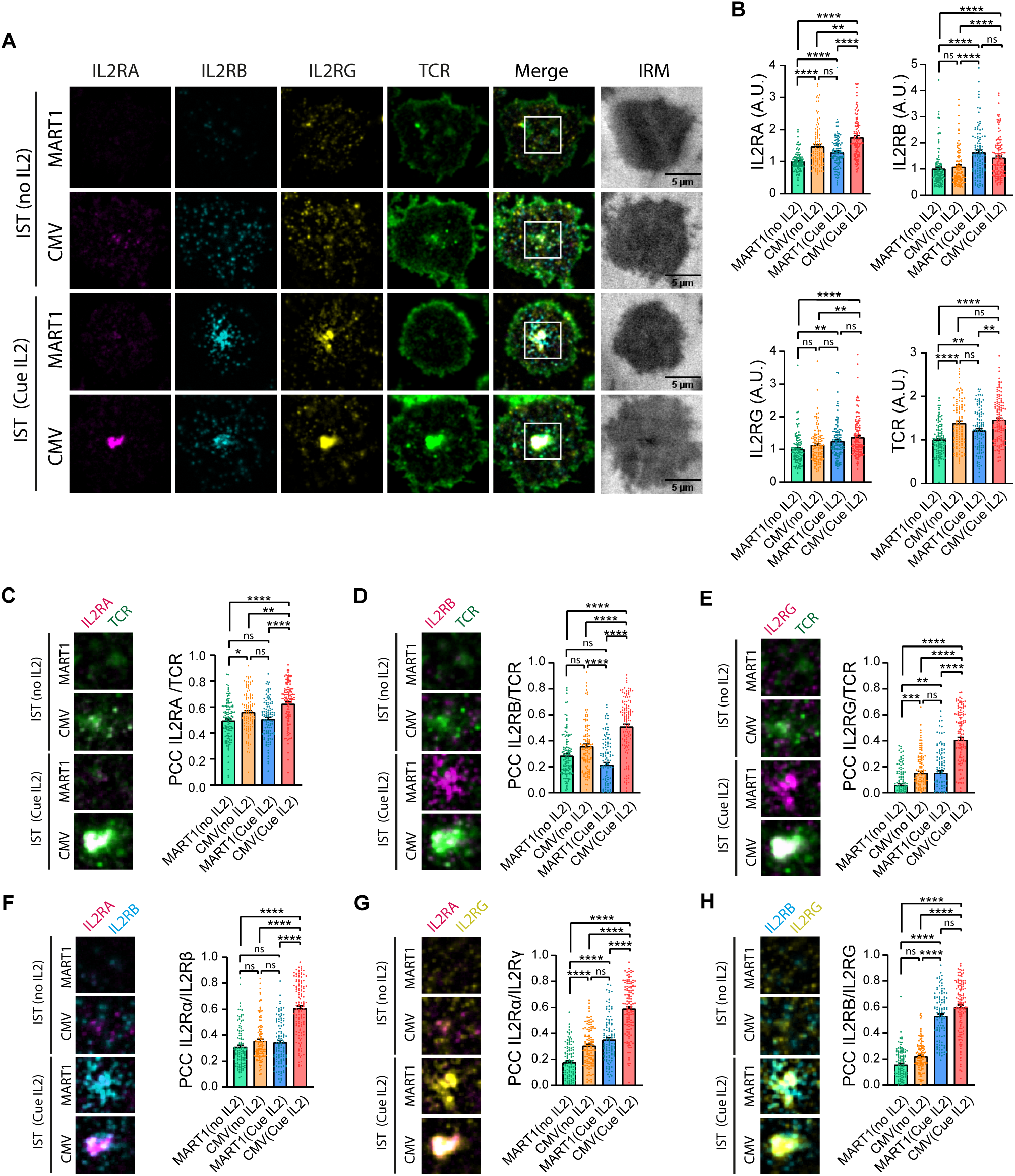
IST induce IL2 receptor assembly and proximity with the TCR at the IS. **(A)** Representative TIRF images of RA14 CD8^+^ blasts forming IS with SLBs containing the indicated IST with ICAM1, CD80 and CD58. Cells were fixed after 15 min of SLB exposure and stained for the IL2 receptor subunits (IL2RA, magenta; IL2RB cyan; IL2RG yellow) and the TCR (CD3, green). **(B)** Quantification of the mean grey values for each protein (IL2RA, IL2RB, IL2RG and TCR) at the IS. Values were normalized to the negative control MART1 (noIL2). Data is the mean +/- SE, n>100 cells from 3 independent donors; **pv<0.01, ****pv<0.0001; Kruskal-Wallis test. **(C-H)** Quantification of the Pearson’s correlation coefficient (PCC) between the TCR and the IL2RA (c), IL2RB (d) and the IL2RG (e) and between the IL2 receptor subunits IL2RA and IL2RB (f), IL2RA and IL2RG (g) and the IL2RB and IL2RG (h). Data is the mean +/- SE, n>100 cells from 3 independent donors; *pv<0.05, **pv<0.01, ****pv<0.0001; Kruskal-Wallis test. Left images show the zoomed in area for the square inset indicated in (a).

Quantification of subunit recruitment and co-localization revealed additional features of the IST system. Recruitment of IL2RB and IL2RG to the IS was driven by CueIL2, with a secondary contribution from the CMV-peptide. Conversely, recruitment of IL2RA and TCR was dominated by the CMV-peptide with a secondary contribution from CueIL2 (Fig. 5A-B). Surprisingly, CueIL2 increased IL2RA recruitment, despite not direct binding. This was supported by increased co-localization of TCR with IL2RA (Fig. 5C), as well as IL2RB (Fig. 5D) and IL2RG (Fig. 5E) in the presence of CueIL2. Furthermore, the combination of CMV-peptide and CueIL2 in ISTs enhanced co-localization of IL2RA with IL2RB (Fig. 5F) and IL2RG (Fig. 5G). IL2RB and IL2RG co-localization increased with CueIL2, regardless of the peptide presented (Fig. 5H).

Presenting the CueIL2 on a separate IST from the CMV-peptide had little effect on the level of IL2R subunit recruitment but reduced the co-localization of IL2R subunits with each other and the TCR (Fig. S5A-D).

When the CueIL2 was substituted by nativeIL2 (with increased IL2RA and IL2RB binding) the only changes were a small increase in IL2RG subunit recruitment (Fig. S6A-B), a slight decrease on IL2RA colocalization with TCR (Fig. S6C) and a slight increase in IL2RA colocalization with IL2RG (Fig. S6D).

While the co-localization data do not demonstrate formation of molecular complexes required for signaling, proximity on this length scale (∼300 nm) would be a prerequisite for such complexes. These results suggest that simultaneous presentation of IL2 and pMHC ligands at the IS potentiates IL2 receptor assembly and proximity to the TCR complex.

### Simultaneous delivery of IL2 and pMHC boosts IL2 receptor and TCR signaling

IL2 receptor signaling (pSTAT5) and TCR signaling (pZAP70) were visualized at the immunological synapse (IS) of RA14 naïve, memory, and blasted CD8+ T cells exposed to SLB with the IST alongside ICAM-1, CD80 and CD58.

In all T cell states, there was a significant and large increase in pSTAT5 and pZAP-70 (Fig. 6) when the IST presented both the CMV-peptide and CueIL2. Surprisingly, the Mart1-peptide and CueIL2 was often second ranked for both pSTAT5 and pZAP-70, suggesting a synergy of the CueIL2 with the co-stimulatory signals on the SLB. Spatial correlation between pSTAT5 and pZAP70 was also highest with CMV-peptide and CueIL2 ISTs, with Mart1-peptide and CueIL2 ISTs ranking second (Fig. 6D, H and L).

**Figure 6.**
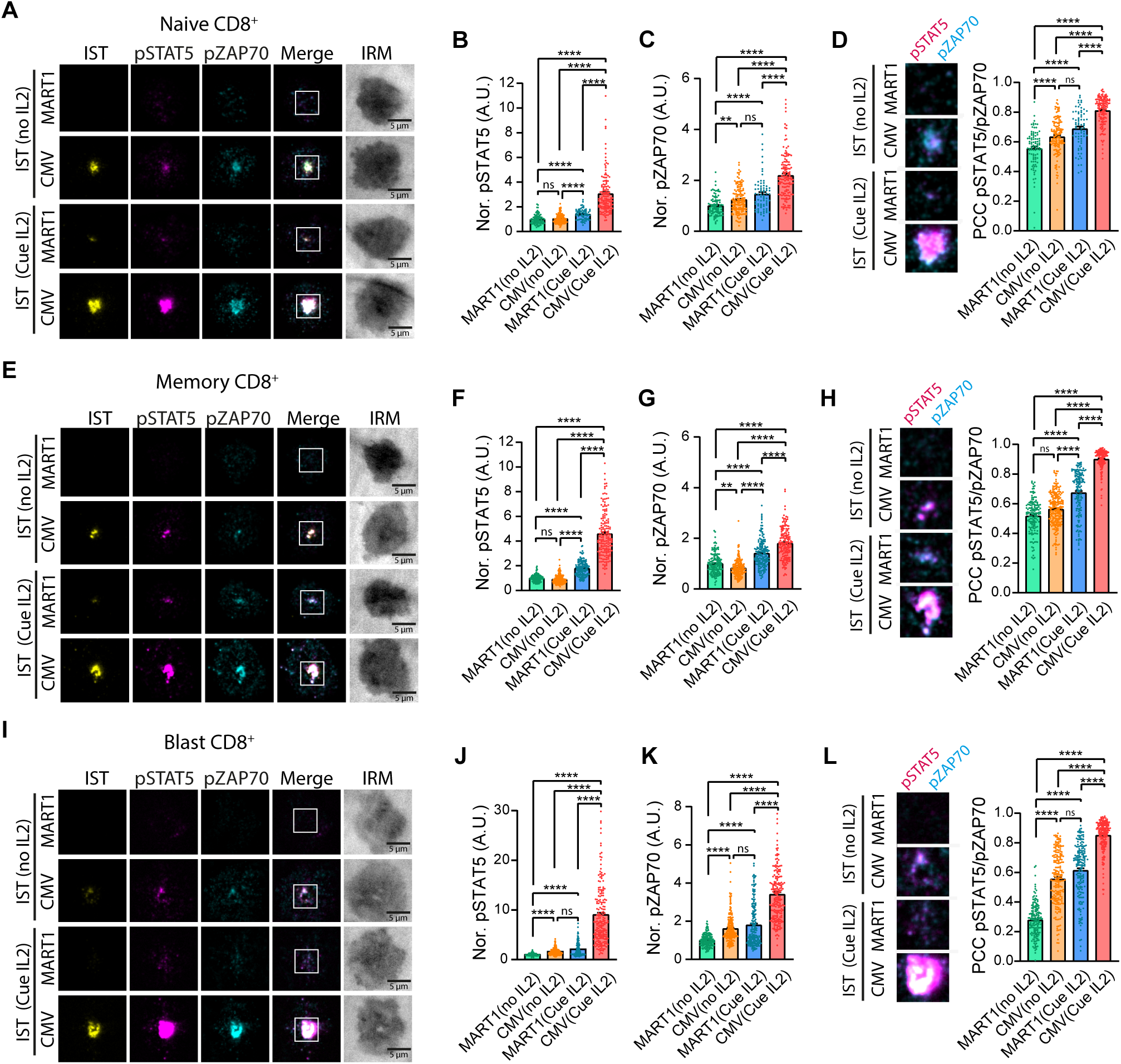
Simultaneous delivery of pMHC and IL2 signals promotes IL2 receptor assembly and TCR proximity to boost signaling. RA14 CD8^+^ Naive **(A-D)**, Memory **(E-H)** and Blasts **(I-L)** T cells forming ISs with SLBs containing the indicated variant of IST AF488 (yellow), ICAM1, CD80 and CD58. Cells were fixed after 15 min of SLB exposure and stained for pSTAT5 (magenta) and pZAP70 (cyan). Middle plots show the quantification of the mean grey values for pSTAT5 and pZAP70 at the IS for Naïve (b, c), Memory (f, g) and Blast (j, k). Values were normalized to the negative control MART1(noIL2). Right plots show the Pearson’s correlation coefficient (PCC) between pSTAT5 and pZAP70 for Naïve (d), Memory (h) and Blast (l). Images in (d, h and l) show the zoomed in area for the square insets indicated in (a, e and i), respectively. Data is the mean +/- SE, n=200 cells from 3 independent donors; **pv<0.01, ****pv<0.0001; Kruskal-Wallis test. CMV, cognate antigen; MART1, negative control antigen.

When CueIL2 was presented separately (CMVnoIL2 + FcCueIL2), pSTAT5 was unaffected, but pZAP- 70 was significantly lower compared to ISTs where both CMV-peptide and CueIL2 were co-presented (Fig. S6E-G). This reciprocal synergy between IL2R and TCR signaling underscores the functional impact of ISTs on enhancing T cell activation. No differences in pSTAT5 or pZAP70 were observed when the IST presented CueIL2 or a Native (wild-type) IL2 (Fig. S7E-G).

### Immuno-STATs act through homotypic T-T cell synapses

To further investigate the mode of action of ISTs in a physiological setting, PBMCs containing RA14 CD8^+^ T cells were incubated with or without 1 µM CMV (CueIL2) ISTs. Next, PBMC samples were stained with antibodies to identify CD4^+^ and CD8^+^ T cells, NK cells, B cells and monocytes and imaged using an ImageStream system (Fig. S8). Treatment with ISTs increased the frequency of doublets (Fig. 7A) and CD3^+^CD8^+^ T-T cell conjugates were the main population enriched by CMV (CueIL2) ISTs (Fig. 7B). In addition, ISTs were visualized at the T-T cell contact, further confirming that these homotypic synapses were triggered by the IST itself (Fig. 7C). IST also accumulated in multiple synapses formed in triplets of CD8^+^ T cells (Fig. 7D). The other main subset of doublets containing ISTs were CD3^+^CD8^+^ T cells – CD14^+^ monocytes. In this case, monocytes engulfed IST molecules together with CD3 molecules from the T cell surface (Fig. 7E-F).

**Figure 7.**
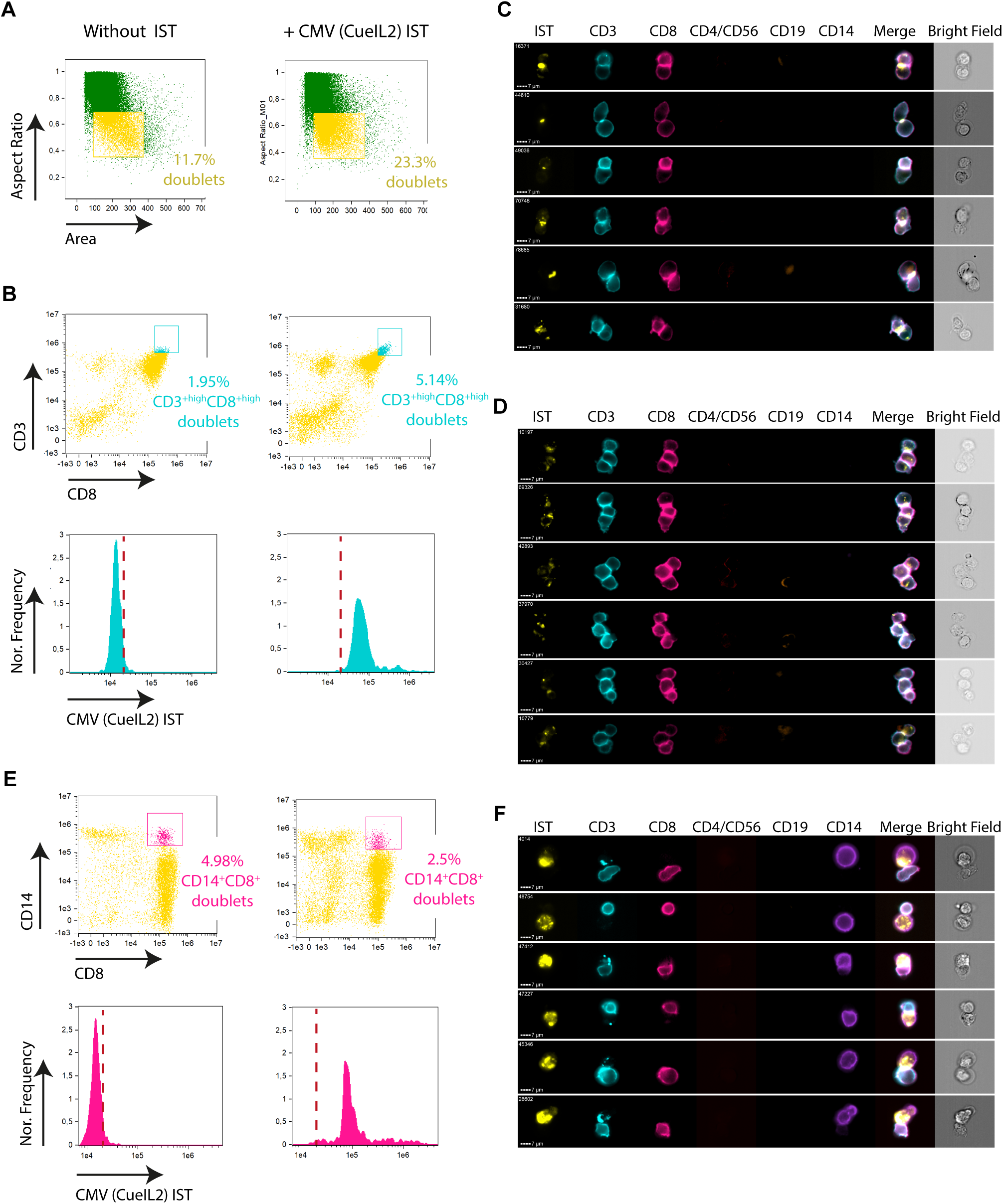
IST function is integrated during homotypic T-T cell immunological synapses. Blast CD8^+^ T cells were electroporated to express the RA14 TCR and reintroduced to the PBMC sample from the same donor. After 2h incubation in a small volume at 37 °C with or without the CMV IL2 IST (1 µM), the cell mix was fixed and blocked (TrueStain, Biolegend) before adding the antibody mix for ImageStream analysis. **(A)** Gating strategy. Images in focus were used to identify the cell doublets (yellow population, left dot plot). **(B)** Doublets were further gated to identify possible CD8^+^ T-T cell conjugates (Dot plots, high values for CD3 and CD8 markers, cyan population). Histograms show the intensity for CMV IL2 IST for the cyan population (possible CD8^+^ T-T cell conjugates). **(C)** Examples of CD8^+^ T-T cell conjugates identified. Note the accumulation of IST at the contact between the two T cells. **(D)** Examples of CD8^+^ T cell multifocal clusters. Note the accumulation of IST at the contact between any two T cells. **(E)** Doublets were further gated to identify possible CD8^+^ T cell conjugates with CD14^+^ monocytes (Dot plots, doublets positive for CD8^+^ and CD14^+^ markers, magenta population). Histograms show the intensity for CMV IL2 IST for the magenta population (possible CD8^+^ T cell – CD14^+^ monocytes conjugates). **(F)** Examples of CD8^+^ T cell – CD14^+^ monocytes conjugates identified. Note the engulfment of ISTs by the CD14^+^ cell together with CD3. Data is representative from two independent donors. The rest of the channels imaged are shown in Fig. S8.

## DISCUSSION

IL2 receptor subunit expression varies across T cell subsets, fine-tuning IL2 sensitivity. Competing interleukin receptor subunits add regulatory complexity. We have confirmed expression levels of these receptor subunits in naïve, memory and blasted CD8^+^ T cells and visualized their expression and distribution at the IS.

While polarization of IL2 release and signaling towards the IS has been described (Sabatos et al., 2008, Reichert et al., 2001, Huse et al., 2006), the precise distribution of the IL2 receptor subunits within the IS was previously unknown, resulting in an incomplete understanding of how IL2 signaling is organized during T cell activation. We found that IL2 receptor subunits display punctate patterns within the IS, and that these clusters can be configured homogenously across the interface, centralized in the IS or in distal ring/corolla . This central accumulation likely involves endocytosis or ectocytosis, as recently described for the TCR (Kvalvaag et al., 2023), and was more frequent in the presence of free IL2 in the media. Previous studies show that IL2 receptor/IL2 complexes are rapidly internalized upon IL2 binding, down-regulating IL2 responses and availability. In a human IL2 dependent T lymphoma cell line, IL2 enters cells by endocytosis in a complex containing all subunits, but IL2RA is recycled back to the plasma membrane, while IL2RB and IL2RG are targeted to late endosomes/lysosomes and degraded (Hemar et al., 1995, Duprez et al., 1994). Imaging of the IL2 receptor subunits by TIRFM revealed some puncta containing all subunits, but the majority were non-overlapping, suggesting that only a small percentage of the IL2 receptor subunits are forming a functional complex, even in the presence of IL2. This may be due to competition with other receptor subunits, such as the case described for the IL7R outcompeting the IL2RB/A for IL2RG at the IS of autophagy-deficient T cells (Zhou et al., 2022) and the ability of IL7 to suppress responses to IL4 and IL21 in naïve T cells due to competition for limited IL2RG (Gonnord et al., 2018).

IL2 and IL2 receptor subunits were also identified in EV released by CD8^+^ T cells. While IL2, IL2RA and IL2RB were enriched in EVs from activated CD8^+^ T cells, IL2RG was enriched in EVs from resting CD8^+^ T cells. EV were mostly single positive for IL2R subunits, suggesting sorting operates on individual subunits post-dissociation. Along these lines, IL2RG contains a PEST sequence that could represent one such sorting sequence (Yu et al., 2000). EVs from activated T cells can induce pSTAT5 in naïve CD8^+^ T cells, indicating a mode of T-T collaboration beyond soluble cytokines.

In addition to characterizing the distribution of the IL2 receptor at the IS, we visualized its activity via STAT5 signaling. Blast cells showed the highest sensitivity to IL2, yet STAT5 signaling was evident at the IS across all T cell states. As previously described for T-T cell contacts, STAT5 signaling was polarized towards the synaptic cleft, mirroring the IL2 receptor complex (Sabatos et al., 2008). Formation of the IS induced pSTAT5 in microclusters overlapping with TCR-mediated pZAP70 microclusters, indicating a spatiotemporal link between TCR and IL2 receptor signaling. TCR microclusters exclude CD45 (Varma et al., 2006), which is a promiscuous transmembrane tyrosine phosphatase that has been shown to target Jak kinases and inhibit cytokine receptor signaling (Irie-Sasaki et al., 2001). This suggests that TCR engagement and IS formation can trigger pSTAT5 signaling. In fact, previous work also shows that pSTAT5 can be directly induced by TCR pathways (Beyer et al., 2011). This regulation is of reciprocal nature, as levels of pZAP70 were also dependent on IL2 activity and previous work shows that IL2 can modulate TCR signaling threshold (Au-Yeung et al., 2017).

To further explore the spatiotemporal relationship between the IL2 and TCR pathways and the therapeutic relevance of crosslinking both signals, we visualized and analysed ISs induced by ISTs. ISTs simultaneously recruited and activated the TCR and the IL2 receptor subunits at the IS.

Remarkably, Ag presentation was enough to trigger the assembly of functional high-affinity IL2 receptors and IL2 presentation boosted TCR signaling to the IS, even in the absence of Ag. Additionally, TCR engagement increased the proximity between the IL2 receptor and the TCR complex itself, further suggesting that the synergy between these pathways relies on their spatial proximity, thereby explaining the maximal activity achieved by ISTs. This suggests that IST act in *cis* mode, meaning that the same IST molecule recruits both IL2 receptor and TCR. Indeed, recent studies highlight the importance of *cis*-targeted IL2 variants in enhancing T cell immunotherapy, although in these instances PD-1 or CD8 receptors were used to localize IL2 to T cells (Codarri Deak et al., 2022, Hashimoto et al., 2022, Quijano-Rubio et al., 2023). In contrast, *cis*-targeting of IL2 through TCR specificity offers superior selectivity, as it engages disease-relevant T cells. In cancer patients, most T cells expressing PD-1 or CD8 are likely not specific to tumour antigens.

Our study suggests that ISTs operate through homotypic T cell-T cell (T-T) and heterotypic T cell-monocyte interactions. Analysis of Ag-specific T cells co-cultured with autologous PBMCs and ISTs showed that the greatest number of IST-induced contacts were T-T. Homotypic T-T interactions occur in lymphoid tissues, probably enhanced by LFA-1/ICAM-1 dependent aggregation (Rothlein and Springer, 1986, Rothlein et al., 1986, Sabatos et al., 2008). We speculate that the IST can bridge TCRs on adjacent T cells specific for the same pMHC. If such IST-driven TCR microclusters were formed, then the attenuated IL2 units could help overcome Treg mediated IL2 scavenging by being protected not only in the close T-T synapse, but within the confines of TCR microclusters. This provision of IL2 may be critical for survival and expansion of tumour reactive-CD8^+^ T cells that may otherwise struggle to find sufficient effector CD4^+^ T cell-derived IL2 in the presence of tumour protective Tregs (Liu et al., 2015).

Presentation of the IST by monocytes is suggested by the engulfment of the IST together with CD3 molecules by monocytes in contact with CD8^+^ T cells. Fc mutations (L234A; L235A) in ISTs attenuate, but do not eliminate common fragment receptor (FcgR) binding (Quayle et al., 2020). T cell stimulation can lead to budding of TCR toward the APC by ectocytosis in both CD4^+^ and CD8^+^ T cells (Kvalvaag et al., 2023), which would account for the appearance of IST and CD3 in the monocytes. Internalized materials from T cells may also include cytotoxic particles (Balint et al., 2020), potentially reducing T- monocyte conjugates in the presence of ISTs. Polarized T cells are resistant to these effects (Li and Orange, 2021, Rudd-Schmidt et al., 2019, Zhang et al., 2006) consistent with the increase in T-T interaction with the IST present.

In vivo immune activation through cytokines or surface receptors in response to antigenic challenge is a highly regulated process. Our data provide key insights into the interplay between TCR and IL2 signaling at the IS. The extension of this biology via ISTs further highlights the importance of TCR engagement in context of IL2 signals. Harnessing the TCR specificity is an elegant approach to selectively activate disease-specific T cells, which allows for a natural mechanism for generation of a therapeutic index for IL2. This contrasts to other approaches where different variants of IL2 are being pursued with little to no consideration towards disease-specific T cells. An extension of the learnings here could be deployed to further our understanding of additional T cell-activating cytokines (IL7, IL15, IL21 etc.) and cell surface receptors (CD28, 41BB etc). To note, many prior clinical approaches in oncology have attempted to harness these potent activation pathways by deploying systemic cytokine administration and/or receptor-agonist antibodies. Most of such studies have had challenges with poor tolerability, toxicities and minimal clinical benefit. Our mechanistic data with ISTs may provide an avenue to selectively drug the anti-tumour T cell repertoire to enhance clinical benefit while preserving patient safety. Indeed, to this end, the emerging clinical data with CUE-101 and CUE-102, which are two different ISTs currently in clinical development of solid cancers, demonstrate clinical efficacy and favourable tolerability (Zhang et al., 2021).

In conclusion, our work demonstrates that *cis*-targeting attenuated IL2 and pMHC molecules, not only effectively overcomes the challenge of selectively delivering IL2 to antigen-specific T cells, but also represents a new strategy to amplify the inherent synergy between these signaling pathways both at the single-cell level and as a modulator of the collective T cell behaviour. The former, by enhancing synaptic TCR and IL2 signaling to levels surpassing the additive effects of each ligand alone. The later, by inducing homotypic T-T cell synapses that leverage cytokine-sharing and quorum-sensing properties, offering a proliferative advantage to low-frequency tumor-specific CD8^+^ T cells. This discovery represents a promising leap forward in precision immunotherapy, with far-reaching implications for targeted cancer treatment.

## MATERIALS AND METHODS

### Isolation of primary human CD8^+^ T cells

Primary human T cells were isolated from leukoreduction system (LRS) chambers, which were supplied by the UK National Health Service Blood and Transplant and sourced from healthy, anonymous donors. The use of these chambers was authorized by the NHS Health Research Authority, National Research Ethics Service (REC reference 11/H0711/7). CD8+ T cells were purified using the negative selection kits RosetteSep™ Human CD8+ T Cell Enrichment Cocktail (StemCell Technologies). To obtain primary CD45RA+CD8+ naïve T cells or CD45RO+CD8+ memory T cells, enriched CD8+ T cells were further purified using EasySep™ Human Naïve CD8+ T Cell or EasySep™ Human Memory CD8+ T Cell Isolation Kits (StemCell Technologies), respectively. Cells were cultured at 37 °C, 5% CO2, in RPMI 1640 medium (Life Technologies) supplemented with 10% FCS, 4 mM L-glutamine, 1% Penicillin-Streptomycin (Gibco), 1% Non-Essential Amino Acids Solution (Thermofisher Scientific), 10mM HEPES (Life Technologies). To generate blast CD8+ T cells, cells were cultured in the presence of anti-human CD3/CD28 Dynabeads (Gibco) at a ratio of one bead per cell and in the presence of 50 U/ml of IL-2. Dynabeads were removed by magnetic separation after 3 days and cells were kept in culture in the presence of IL-2 until day 7.

### Expression plasmids and mRNA electroporation

The sequences for the CMV ppp65-specific RA14 TCR were provided by Cue Biopharma. RA14 TCRA and TCRB constructs were synthesised using GeneArt services from ThermoFisher and subcloned into a pGEM-GFP-64A plasmid between HindIII and KpnI. A Kozak sequence was added and codon optimization for human expression was applied. Both RA14 TCRA and TCRB constructs included a cysteine modification, as previously described in (Abu-Shah et al., 2019). For mRNA synthesis, plasmids were linearized and in vitro RNA transcription and in vitro poly-adenylation were performed using the mMESSAGE mMACHINE T7 ULTRA Transcription kit (Thermofisher Scientific), following manufacturer’s instructions. For electroporation, cells were washed three times in Opti-MEM media (LifeTechnologies) and resuspended at 2-2.5x10^6^ cells in 100 μl and mixed with 5 µg of each mRNA stock (RA14 TCRA, RA14TCRB and CD3ζ). Samples were then transferred to a Gene Pulser/Micropulser electroporation cuvette, 0.2 cm gap (BIO-RAD) and pulsed at 300 V for 2 ms in an ECM 830 Square Wave Electroporation System (BTX). Cells were immediately transferred to supplemented RPMI and cultured at 37 °C, 5% CO2 before usage after 18-24 h.

### Preparation of Supported Lipid Bilayer (SLB) and immunocytochemistry

To prepare SLBS, glass coverslips (Nexterion) were plasma cleaned and mounted onto six-channel chambers (Ibidi). Small unilamellar liposomes were prepared using 1,2-dioleoyl-sn-glycero-3- phosphocholine (Avanti Polar Lipids Inc.) supplemented with 12.5% 1,2-dioleoyl-sn-glycero-3-[(N-(5- amino-1-carboxypentyl) iminodiacetic acid) succinyl]-Ni (Avanti Polar Lipids Inc.). Channels in Ibidi chamber were covered with liposome mixture, blocked and washed. SLB were then incubated with the indicated mix of His-tagged proteins to achieve the desired density of molecules on the SLB: UCHT1 (30 molecules/ μm^2^), ICAM1 (200 molecules/μm^2^), CD80 (200 molecules/μm^2^), CD58 (200 molecules/μm^2^), pMHC (100 molecules/μm^2^). His-tagged proteins were produced in house and, when required, conjugated with the AlexaFluor (AF) dyes of interest (405, 488, 568 or 647). For experiments with ISTs, SLBs were incubated with 200 nM of His-tag D-Peptide derived from Protein A (Bankpeptide, Hefei, China) (Zhang et al., 2013) for 20 min and then the protein mix was added with the indicated IST variant at 3 nM. To generate the ISs, cells were exposed to the bilayers at 37 °C for the indicated amount of time min, fixed with 4% PFA and washed. For immunocytochemistry, cells were permeabilized, blocked and incubated with the primary antibody of interest (anti-pSTAT5 AF647, BD Biosciences; anti-pZAP70 CST; anti-pZAP70 PE BD Biosciences; anti-IL2RA FITC clone 2A3 BD Biosciences, anti-IL2RB APC clone Mik-β3 (RUO), BD Biosciences; anti-IL2RG, ThermoFisher; anti-CD3 PE, BioLegend;) for 1h at room temperature (RT) or overnight at 4°C. When required, samples were further incubated with an AF-conjugated secondary antibody for 1h at RT.

### TIRFM, Airyscan confocal microscopy, Opera Phoenix high content maging and image analysis

TIRF imaging was performed on an Olympus IX83 inverted microscope with a TIRF module. The instrument was equipped with an Olympus UApON 150x TIRF N.A 1.45 objective, 405 nm, 488 nm, 568 nm and 640 nm laser lines and Photomertrics Evolve delta EMCCD camera. For live imaging experiments, SLBs were transferred to a pre-heated incubator on top of the TIRF microscope and cells were added in the well. Airyscan imaging was performed using a Zeiss 980 LSM using a 63x oil 1.40 NA objective. Images were visualized using ZenBlue Software or Fiji (ImageJ). For High Throughput Imaging, SLBs were prepared on 96-well plates and imaged with the opera Phoenix High Content Imaging System. Imaging was performed at RT. Image analysis was performed using Fiji (ImageJ). Pearson correlation coefficients (PCCs) were calculated using the JACoP plugin. To quantify the mean grey value per cell, synaptic contacts were segmented using the IRM image.

### Intracellular Flow Cytometry

To assess levels of intracellular phosphorylated proteins, cells were fixed in 2% PFA for 15 min, washed and gently resuspended in ice-cold 90% methanol and incubated for a minimum of 3h at -20°C. Next, cells were washed and incubated with the corresponding antibody in 0.5% BSA PBS solution for 1h. Samples were washed and analysed using the LSR II Flow Cytometer (BD Biosciences). Data were analyzed using FlowJo version 10.7.1.

### Single-cell RNA sequencing (scRNA-seq)

The raw scRNA-seq data was downloaded from Gene Expression omnibus (GEO) under accession number GSE126030. The article corresponding to the repository illustrates that all the T cells were obtained from human PBMCs. These T cells were activated in the presence of anti-CD3/anti-CD28 beads for 48 hours and compared to the resting T cells which were cultured in media alone. The fastq files obtained from GEO were processed using the Cell Ranger (v3.1.0) counts pipeline. The sequencing reads were aligned to Human GRCh38-3.0.0 and the cell count matrices were obtained. These filtered gene–cell matrices were then read into the Seurat v4.0 pipeline implemented in R for downstream analysis. We followed the standard Seurat pre-processing workflow with QC based cell filtration, data normalization and scaling. We further identify major cell types with highly variable genes and performed Jackstraw analysis to isolate significant principal components. The UMAP embeddings were then computed using the isolated PCs and known marker genes were used to characterize cell types.

### Isolation and purification of extracellular vesicles and proteomic analysis

Cells were incubated in serum free media for 48 h followed by collection of media. The media supernatant was spun at 300g to remove cells followed by a 4,000g centrifugation to discard cell debris. The supernatant was concentrated using 100 kDa molecular weight cut-off Amicon centrifugal filter units (Merck Millipore, Billerica, Massachusetts, USA) to 2 ml and subsequently loaded on a Sepharose 4 Fast Flow resin column (10 mm x 300 mm; GE Healthcare, Little Chalfont, UK), connected to the ÄKTA pure system (GE Healthcare) and eluted at 0.5 ml/min flow rate using PBS as the eluent. Chromatogram was recorded using absorbance at 280 nm. 0.5 ml fractions were collected, and EV-containing fractions were pooled and concentrated to 2 ml with 10 kDa molecular weight cut-off Amicon centrifugal filter units (Merck Millipore). For the proteomic analysis, EV samples (750 μL) were reduced with 5 μL of 10 mM tris(2-carboxyethyl)phosphine and alkylated with 50 mM of iodoacetamide for 30 min each, then acidified with 12% phosphoric acid 10:1 vol:vol, and then transferred to S-trap columns. Then, samples were precipitated using 1:8 vol:vol dilution of each sample in 90% methanol in 100 mM Triethylamonium bicarbonate and digested with trypsin (Promega, #V1115) overnight at 37° C. The samples were then run on a LC-MS system comprised of an Evosep One and Bruker timsTOF Pro. For proteomic analyses the raw files were searched against the reviewed Uniprot Homo sapiens database (retrieved 2,01,80,131) using MaxQuant version 1.6.10.43. The intensities were baselined using NTA concentration and average-based normalization was carried out to obtain the enriched set of proteins in each EV sample.

### NanoFlow Cytometry

The Flow NanoAnalyzer (NanoFCM Co., LTD) was used to perform Nano flow cytometry analysis according to the manufacturer’s instructions. To construct a calibration curve and enable the conversion of side scatter intensities to particle size, a Silica Nanospheres Cocktail (S16MExo, NanoFCM) was employed as the size standard. In addition, a concentration standard (200nm PS QC beads, NanoFCM) was used to calculate particle concentrations. A 488nm laser at 25/40 mW, 10% ss decay was utilized, and the detectors were equipped with 525/40 (AF488) and 580/40 (PE) bandpass filters. Antibodies were allowed to bind for 30 minutes on ice, followed by removal of unbound antibodies through centrifugation for 1 hour at 100,000g at +4°C. Labelled vesicles were then resuspended in 50 µL of PBS. Sample and label specific signals were defined using buffer alone (PBS), unstained samples, samples stained with isotype controls, and auto-thresholding.

### Multispectral flow cytometry (ImageStream) assay

CD8^+^ T cells were purified from a PBMC sample. Some PBMCs were kept frozen while purified CD8^+^ T cells were blasted as usual. On day 7, blast CD8^+^ T cells were electroporated with the RA14 TCR mRNA 24h prior to coculture experiment. For co-culture, 2 million RA14 CD8^+^ T cells were added on 4 million of freshly thawed PBMCs and incubated for 2h at 37°C in a small volume to favour cell contacts. When indicated, CMV (CueIL2) IST 1µM was added to the media. Next, cells were fixed with 2% PFA for 10 min, washed and blocked with TrueStain buffer (Biolegend) for 10 min. Finally, samples were stained with the antibody cocktail for ImageStream analysis (UCHT1-PE, Biolegend; CD19- AF594, Biolegend; CD14-BV421, Biolegend; CD8-AF700, Biolegend; CD4-APC/Fire750, Biolegend; CD56-APC/Fire750, Biolegend). Data were acquired on an ImageStream MK II (Amnis) with INSPIRE V4 software and analyzed using IDEAS 6.2 software. Briefly, in-focus images were used to gate doublets based on the area and the aspect Ratio of the bright field images. Subsequent gating was carried out using the specific marker staining. High CD3^+^CD8^+^ were considered possible T-T cell doublets.

### Statistical analysis

All statistical tests were performed using GraphPad Prism 9 software. Mann-Whitney test and Kruskal-Wallis test were used for statistical analysis and p<0.05 was considered statistically significant.

## DATA AVAILABILITY

The data are available from the corresponding author upon reasonable request.

## ACKNOWLEDGEMENTS

The authors thank MJA Wood for the use of the Nano Flow Cytometer, as well as the essential support from E. Kurz. We thank J Raybin, W Bautista, J Soriano, F Zhao, A Vakkasoglu, J Ross, S Low, and S Cemerski of Cue Biopharma for providing reagents, methods, and input. We would also like to thank all the anonymized blood donors who contributed to our study. This work was supported by Cue Biopharma, the Kennedy Trust for Rheumatology Research Cell Dynamics Program (KENN202117) and the Wellcome Trust (100262Z/12/Z and 224343/Z/21/Z). JC was supported by Cue Biopharma postdoctoral fellowship and a Cancer Research Institute Irvington Fellowship (CRI4503). AJK is supported by the Clarendon Fund, and the Interdisciplinary Bioscience DTP at the University of Oxford (UKRI–BBSRC grant no. BB/M011224/1). SV and LC were supported by the European commission Horizon 2020 (ERC-2021-SyG_951329). JC and MLD received research funding from Cue Biopharma. SNQ, RJM and AS are employees of Cue Biopharma. AJK and MLD are co-founders of GranzaBio. The authors have no additional financial interests.

