## Supplementary Material for "Synaptic synergy of T cell receptor and interleukin 2 receptor in CD8^+^ T cells"

### SUPPLEMENTARY FIGURE LEGENDS

**Figure S1:** *IL2 redistributes the IL2* receptor *complex towards the centre of the IS.* RA14 CD8$^+$ T cells were exposed to Ag-specific SLBs in the presence or absence of 50 U/ml of IL2 to form ISs and fixed after 10 min. Samples were then permeabilized and immunostained against the IL2 receptor subunits. **(A)** Schematic illustration of the classification criteria to classify the IL2 receptor distribution into homogeneous (left), central (middle) or annular (right). **(B-D)** Representative TIRFM images of the different phenotypes (Homogeneous, Central and Annular) observed for the IL2RA, IL2RB and IL2RG for each subset of CD8$^+$ T cells; (b) blast, (c) memory and (d) naïve. Histograms show the frequency for each phenotype as quantified by visual inspection for the no IL2 and 50 U/ml IL2 conditions. Data is the mean of three independent donors with >50 cells/donor.

**Figure S2:** *IL2 modulates the abundance of the IL2* receptor *complex at the IS.* Blast, Memory and Naïve RA14 CD8$^+$ T cells were exposed to antigenic pMHC SLBs to form ISs and fixed after 10 min. Samples were then permeabilized and immunostained against the IL2 receptor subunits. The synaptic contact was imaged with TIRFM. Representative images are shown in Fig. 1g. **(A)** Dot plots show the quantification of mean intensity for IL2RG, IL2RB and IL2RA at the IS in the presence of 50 U/ml of IL2. Histogram shows the mean +/- SE, n>55 cells. ns, not significant, **pv<0.01, ****pv<0.0001; Mann-Whitney test. **(B-D)** Synapses were let to form by Blast (b), Memory (c) or Naïve (d) RA14 CD8$^+$ T cells in the presence or absence of 50 U/ml of IL2 for 10 min before fixation. Violin plots show the quantification of the mean intensity values for each IL2 receptor subunit at the IS normalized to the condition without IL2 (No IL2). Data shows the median and quartiles of >150 cells, from 3 independent donors. ns, not significant; *pv<0.05; **pv<0.01; ***pv<0.001; Mann-Whitney test.

**Figure S3:** *Synaptic assembly of the IL2 receptor complex.* Blast, Memory and Naïve RA14 CD8$^+$ T cells were exposed to antigenic pMHC SLBs to form ISs and fixed after 10 min. Samples were then permeabilized and immunostained against the IL2 receptor subunits. The synaptic contact was imaged by TIRFM. **(A)** Representative TIRF images of a RA14 CD8$^+$ blast T cell forming a synaptic contact with the SLB. Each IL2 receptor subunit image (input image; IL2RG, yellow; IL2RB, cyan; IL2RA, magenta) was segmented into a binary mask to detect the subunit clusters. Further masks were created from the intersection ($\cap$) between the masks of each pair of subunits (double mask). The interference reflection microscopy (IRM) image was used identify the whole cell contact. Next, the mean intensity value for all the subunits was measured within each mask. **(B)** Percentage of the total IL2RG intensity (within the IRM area) that is within the IL2RA mask. **(C)** Percentage of the total IL2RA intensity (within the IRM area) that is within the IL2RB (left plot), IL2RG (middle plot) or the intersection between the IL2RG and IL2RB mask (right plot). **(D)** Percentage of the total IL2RB intensity (within the IRM area) that is within the IL2RG (left plot), IL2RA (middle plot) or the intersection between the IL2RG and IL2RA mask (right plot). **(E)** Quantification of the Pearson's Correlation Coefficient (PCC) for each pair of IL2 receptor subunit at the IS. Representative images in Fig. 1g. Dots show the distribution of the values for >100 cells, from three independent donors. Histogram shows the mean +/- SE. *pv<0.05, ***pv<0.001, ****pv<0.0001; Kruskal-Wallis test.

**Figure S4:** *IL2 effects on synapse formation.* Blast, Memory and Naïve RA14 CD8$^+$ T cells were exposed to antigenic pMHC SLBs to form ISs. SLBs also contained His-tagged ICAM1-AF405, CD80-AF568, and CD58-AF647. When indicated, IL2 was simultaneously added. Samples were fixed after 15 min of SLB and IL2 exposure and immunostained against the TCR (anti-CD3 AF488). **(A-C)** Samples were prepared in a 96-multi well plate containing the SLBs and incubated with increasing concentrations of IL2 in the media, simultaneously to SLB exposure. Synapses were then imaged using an Opera Phenix high content confocal system. Images on the left are representative examples. Bright field images (not shown) were used to segment the cells. Right plots show the mean intensities (MGV) for the TCR (green), ICAM1 (cyan), CD80 (yellow) and CD58 (magenta) normalized to the control (grey line, no IL2 in the media). Data is the mean +/- SE of approximately 250 cells/condition. *pv<0.05, **pv<0.01, ***pv<0.001, ****pv<0.0001; Kruskal-Wallis test (significant differences compared to the control without IL2). **(D-F)** Samples were also prepared for TIRFM imaging. Cells were fixed after 15 min of SLB exposure in the presence or absence of 50 U/ml of IL2 in the media. Images on the left are representative examples. Dot plots on the right show the quantification of the mean intensity values (MGV) at the synapse for the indicated proteins (TCR, green; ICAM1, cyan; CD80, yellow; and CD58, magenta) normalized to the control (no IL2). Data is the mean +/- SE of >150 cells. *p<0.05; ****p<0.0001. Mann-Whitney test.

**Figure S5:** *IL2 effects on IS contact area.* **(A)** *Naïve,* **(B)** Memory and **(C)** Blast RA14 $CD8^+$ T cells were exposed to antigenic pMHC SLBs to form ISs. When indicated, 50 U/ml of IL2 were simultaneously added. Samples were fixed after 10-15 min of SLB and IL2 exposure and imaged with IRM. The image was used to segment the cell contact with the SLB (IS area) and the area of the contact was measured. Data is the mean +/- SE of > 400 cells. *p<0.05; ****p<0.0001. Mann-Whitney test.

**Figure S6.** *Molecular link between IL2 and pMHC signals promotes IL2* receptor *assembly and TCR proximity and boosts signaling of both receptors.* **(A)** Representative TIRF images of RA14 $CD8^+$ Blast T cell forming a IS with a SLB containing cognate ISTs (CMV), ICAM-1, CD80 and CD58. The CueIL2 molecule was part of the IST structure (CMV CueIL2) or presented as a separate Fc molecule (FcCueIL2) together with an IST without IL2 (CMV noIL2). Cells were fixed at 15 min after SLB exposure and immunolabelled against the three subunits of the IL2 receptor (IL2RA, magenta; IL2RB, cyan; IL2RG yellow) and the TCR (green). **(B)** Quantification of the relative mean grey value for the IL2 receptor subunits and the TCR at the IS. Data is normalized to the control (MART1 noIL2, data shown in Fig5). Data is the mean +/- SE, n=125 cells from 3 independent donors; **pv<0.01; ns, not significant; Mann-Whitney test. **(C)** Pearson's correlation coefficient (PCC) between the IL2 receptor subunits and the TCR. Images show the zoomed in square inset indicated in (a). Data is the mean +/- SE, n=125 cells from 3 independent donors; **pv<0.01, ****pv<0.0001; Mann-Whitney test. **(D)** Pearson's correlation coefficient (PCC) between each pair of IL2 receptor subunit. Images show the zoomed in square inset indicated in (a). Data is the mean +/- SE, n=125 cells from 3 independent donors; ****pv<0.0001; Mann-Whitney test. **(E)** Representative TIRF images of RA14 $CD8^+$ Blast T cells forming an IS with a SLB containing cognate ISTs AF488 (CMV, in yellow), ICAM-1, CD80 and CD58 and, when indicated, the FcCueIL2 AF568 (red). Cells were fixed at 15 min after SLB exposure and stained against pSTAT5 (magenta) and pZAP70 (cyan). **(F)** Quantification of the mean grey value for pSTAT5 and pZAP70 at the IS. Data is normalized to the control (MART1 noIL2, data shown in Fig6). Data is the mean +/- SE, n=200 cells from 3 independent donors; ****pv<0.0001; ns, not significant; Mann-Whitney test. **(G)** Pearson's correlation coefficient (PCC) between pSTAT5 and pZAP70. Data is the mean +/- SE, n=200 cells from 3 independent donors; **pv<0.01, ****pv<0.0001; Mann-Whitney test. Images show the zoomed in for the square inset indicated in (e).

**Figure S7.** *IL2/IL2RA binding is not required for IL2 receptor assembly and activity at the IS.* **(A)** Representative TIRF images of RA14 $CD8^+$ Blast T cells forming an IS with a SLB containing cognate ISTs (CMV) with ICAM-1, CD80 and CD58. The IST molecule contained either CueIL2 or NativeIL2.

Cells were fixed at 15 min after SLB exposure and stained against the three subunits of the IL2 receptor (IL2RA, magenta; IL2RB, cyan; IL2RG yellow) and the TCR (green). **(B)** Quantification of the mean grey value for the IL2 receptor subunits and the TCR at the IS normalized to the control (MART1 noIL2, data shown in Fig5). Data is the mean +/- SE, n=125 cells from 3 independent donors; *pv<0.05, **pv<0.01; Mann-Whitney test. **(C)** Pearson's correlation coefficient (PCC) between the IL2 receptor subunits and the TCR and **(D)** between each pair of IL2 receptor subunits. Data is the mean +/- SE, n=125 cells from 3 independent donors; *pv<0.05; Mann-Whitney test. **(E)** Representative TIRF images of RA14 $CD8^+$ Blast T cells forming a IS with a SLB containing cognate ISTs (CMV, yellow). The IST molecule contained either CueIL2 or NativeIL2. Cells were fixed at 15 min after SLB exposure and stained against pSTAT5 (magenta) and pZAP70 (cyan). **(F)** Quantification of the mean grey value for pSTAT5 and pZAP70 at the IS normalized to the negative control (MART1 noIL2, data shown in Fig5). Data is the mean +/- SE, n=200 cells from 3 independent donors; Mann-Whitney test. **(G)** Pearson's correlation coefficient (PCC) between the indicated IL2 receptor subunits and the TCR. Data is the mean +/- SE, n=200 cells from 3 independent donors; **pv<0.01; Mann-Whitney test.

**Figure S8.** Representative images of cell singlets stained with the different surface markers used for ImageStream analysis.

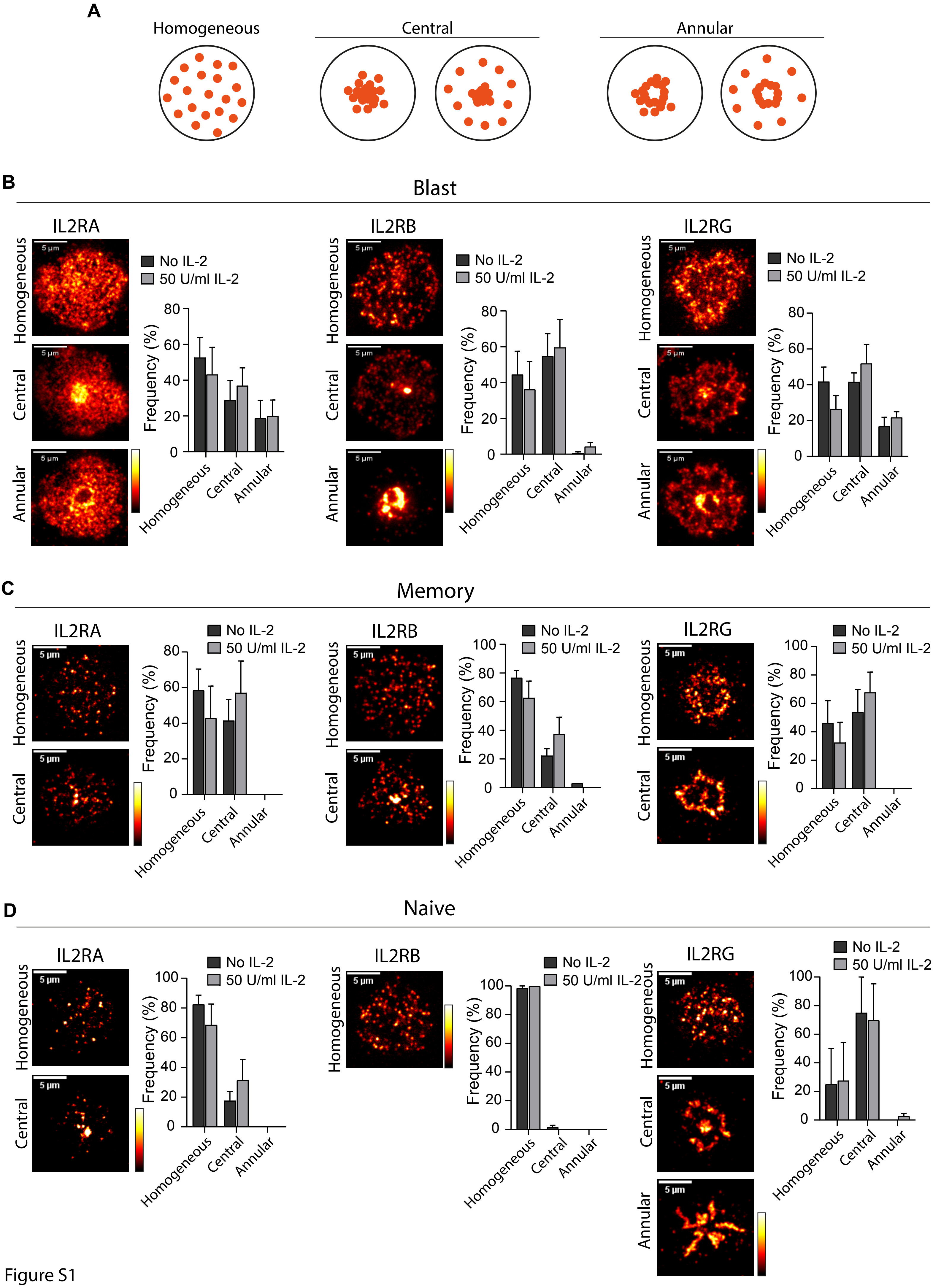
A
Homogeneous
Central
Annular
B
Blast
IL2RA
IL2RB
IL2RG
Homogeneous
Central
Annular
5 µm
No IL-2
50 U/ml IL-2
Frequency (%)
C
Memory
D
Naive

Figure S1

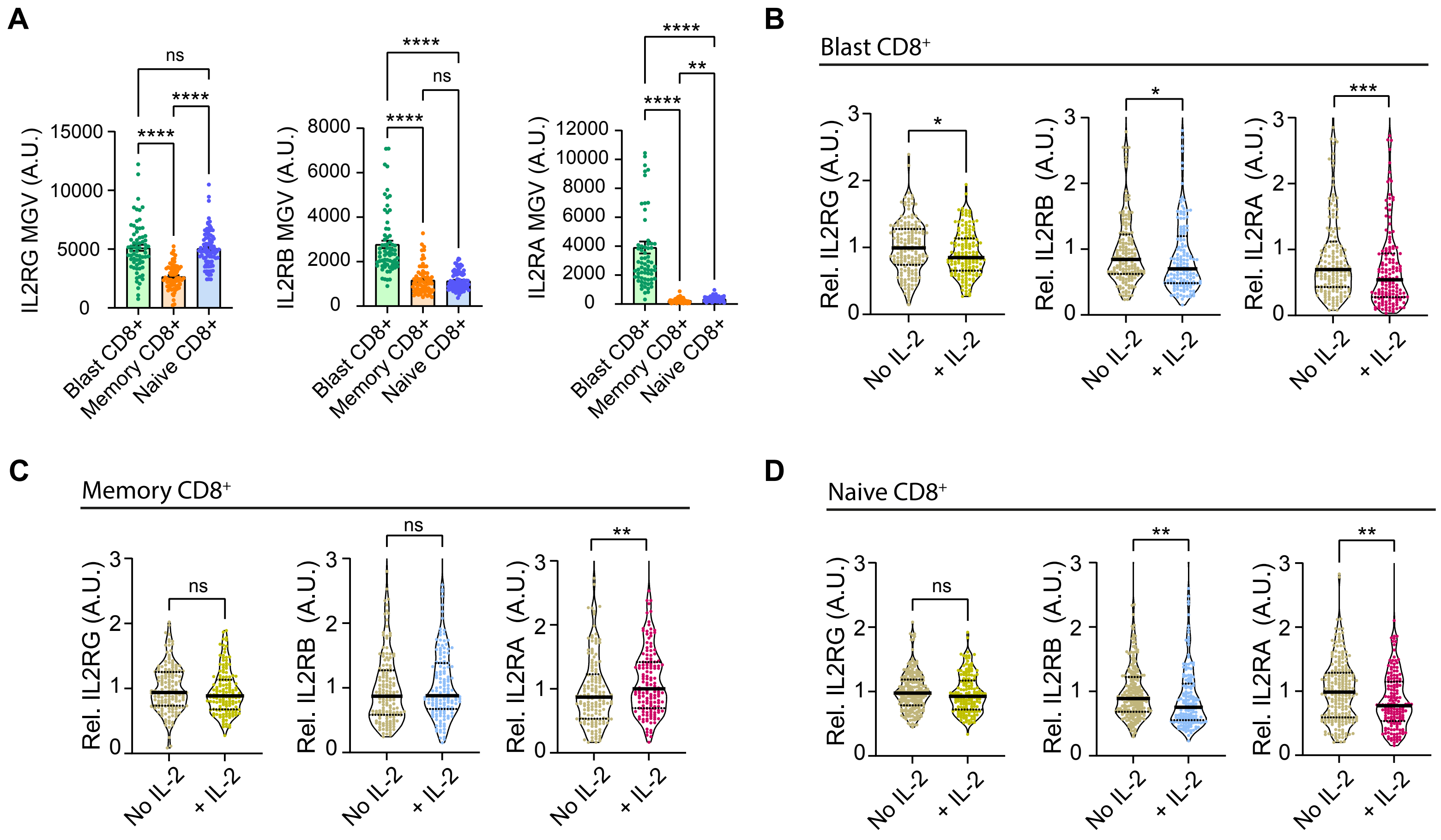
A
IL2RG MGV (A.U.)
IL2RB MGV (A.U.)
IL2RA MGV (A.U.)
Blast CD8+
Memory CD8+
Naive CD8+
B
Blast CD8+
C
Memory CD8+
D
Naive CD8+
Rel. IL2RG (A.U.)
Rel. IL2RB (A.U.)
Rel. IL2RA (A.U.)
No IL-2
+ IL-2
ns
****
***
**
*

Figure S2

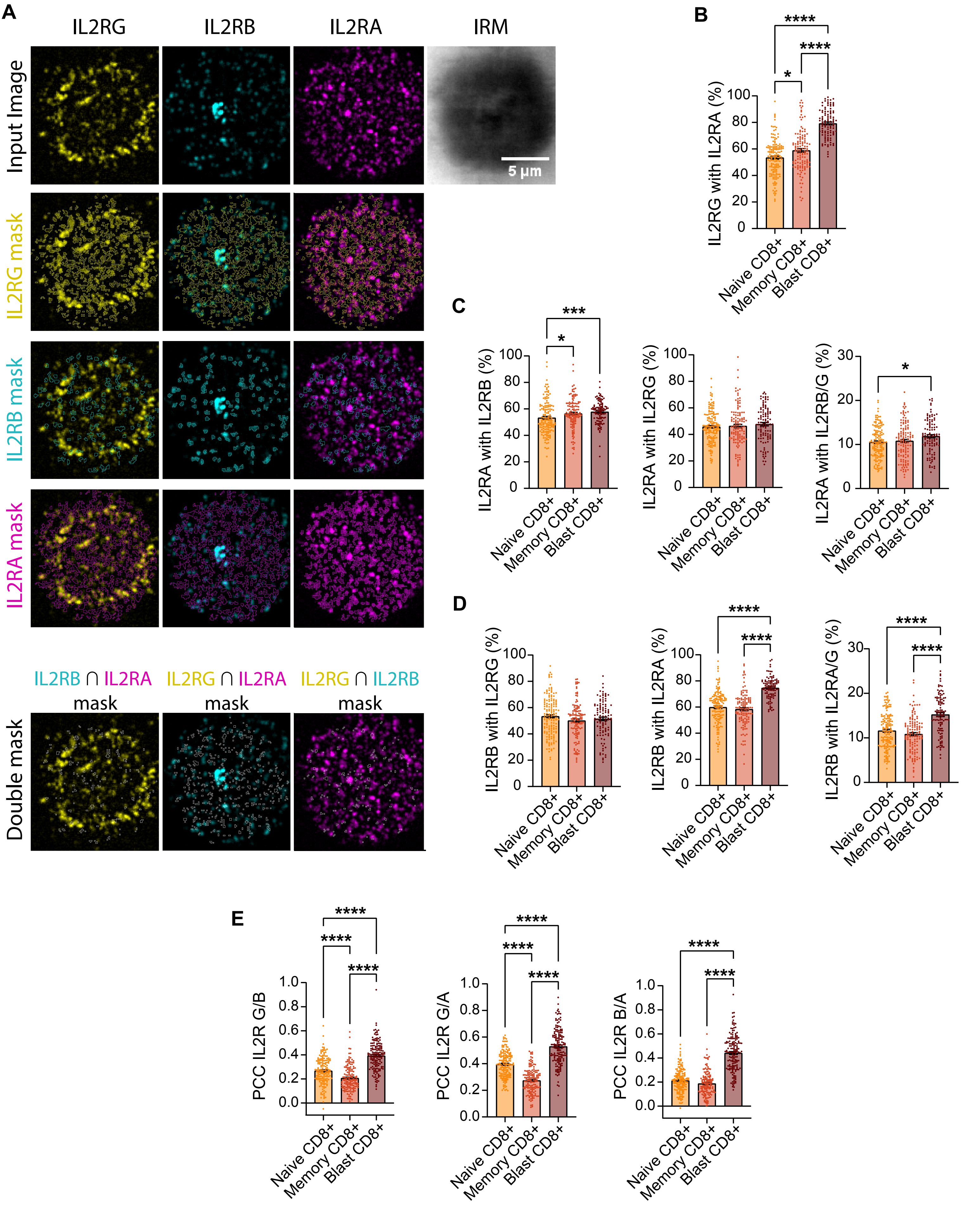
A
IL2RG
IL2RB
IL2RA
IRM
Input Image
5 µm
IL2RG mask
IL2RB mask
IL2RA mask
IL2RB ∩ IL2RA mask
IL2RG ∩ IL2RA mask
IL2RG ∩ IL2RB mask
Double mask
B
IL2RG with IL2RA (%)
Naive CD8+
Memory CD8+
Blast CD8+
C
IL2RA with IL2RB (%)
IL2RA with IL2RG (%)
IL2RA with IL2RB/G (%)
D
IL2RB with IL2RG (%)
IL2RB with IL2RA (%)
IL2RB with IL2RA/G (%)
E
PCC IL2R G/B
PCC IL2R G/A
PCC IL2R B/A

Figure S3

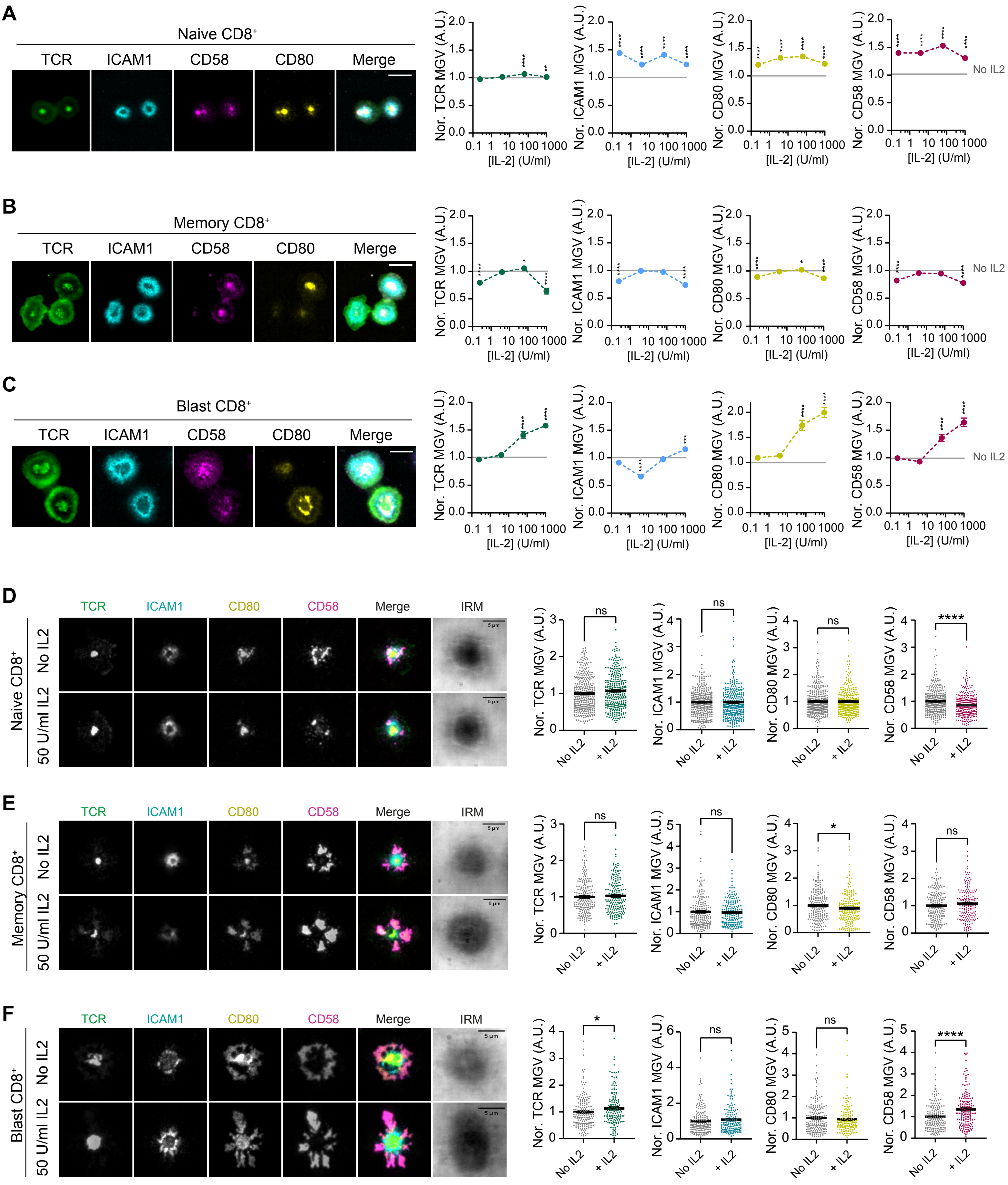
A
Naive CD8⁺
TCR
ICAM1
CD58
CD80
Merge
Nor. TCR MGV (A.U.)
Nor. ICAM1 MGV (A.U.)
Nor. CD80 MGV (A.U.)
Nor. CD58 MGV (A.U.)
[IL-2] (U/ml)
No IL2
B
Memory CD8⁺
C
Blast CD8⁺
D
Naive CD8⁺
No IL2
50 U/ml IL2
IRM
5 µm
ns
****
No IL2
+ IL2
E
Memory CD8⁺
*
F
Blast CD8⁺

Figure S4

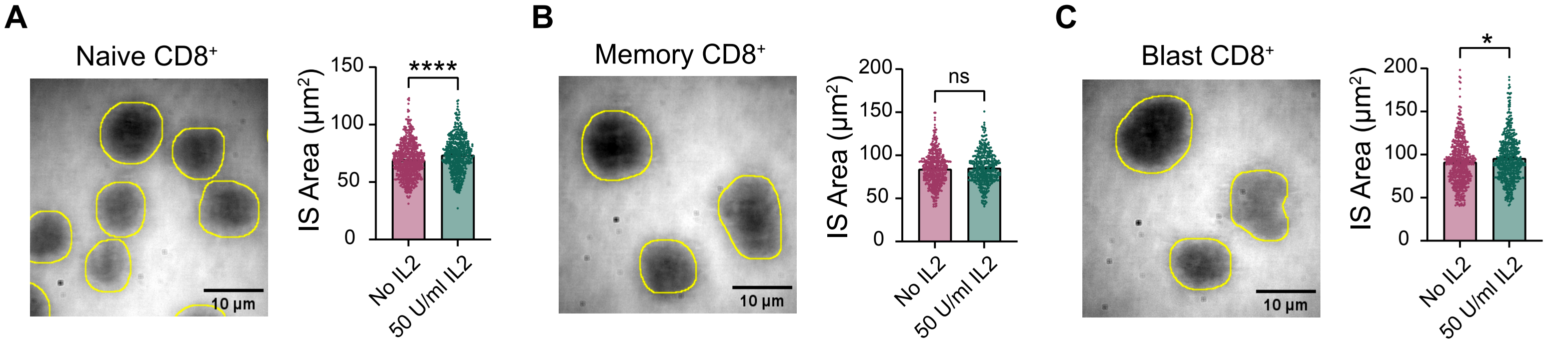
A
Naive CD8⁺
10 µm
IS Area (µm²)
150
100
50
0
****
No IL2
50 U/ml IL2
B
Memory CD8⁺
10 µm
IS Area (µm²)
200
150
100
50
0
ns
No IL2
50 U/ml IL2
C
Blast CD8⁺
10 µm
IS Area (µm²)
200
150
100
50
0
*
No IL2
50 U/ml IL2

Figure S5

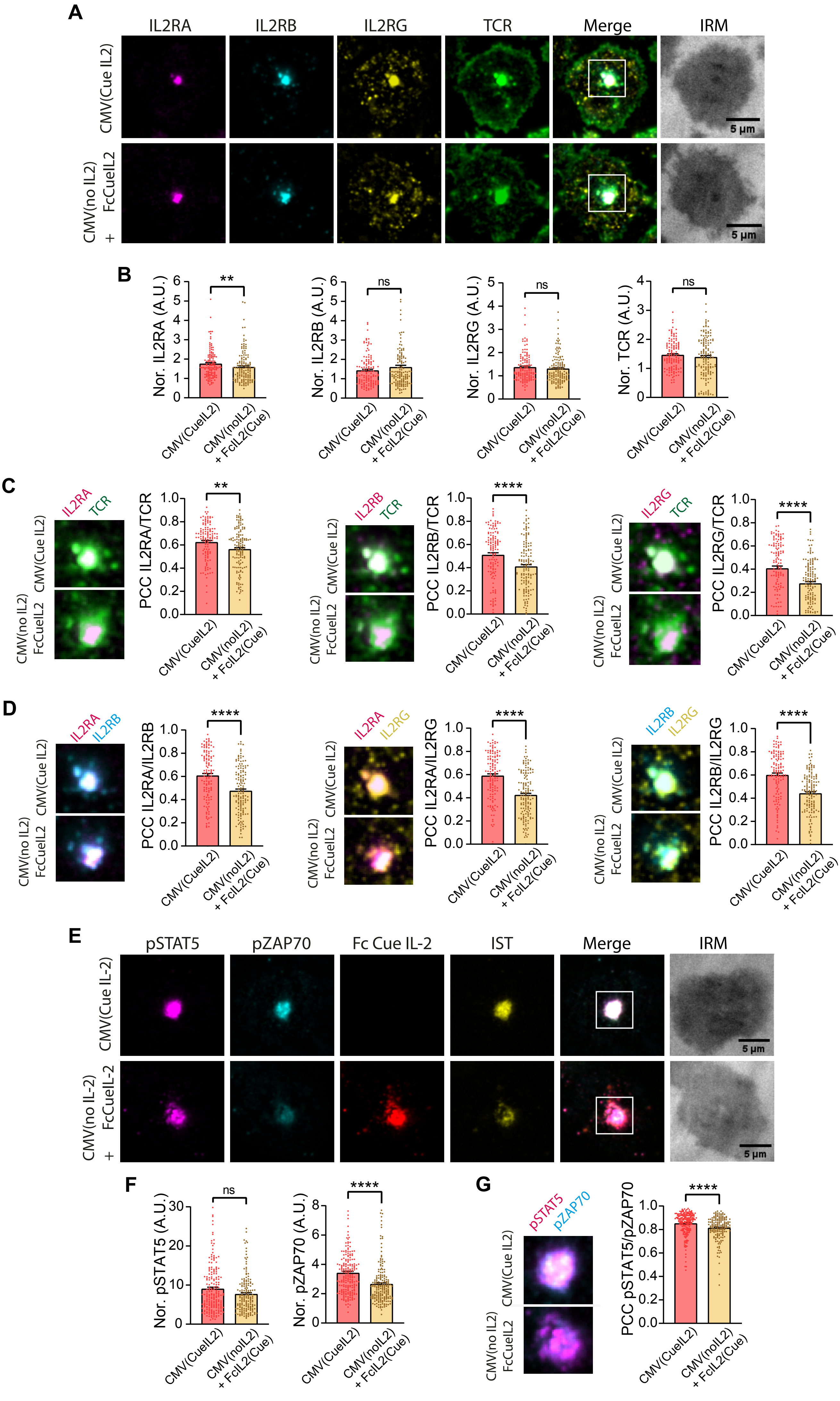
A
IL2RA
IL2RB
IL2RG
TCR
Merge
IRM
CMV(Cue IL2)
CMV(no IL2) + FcCueIL2
5 µm
B
Nor. IL2RA (A.U.)
Nor. IL2RB (A.U.)
Nor. IL2RG (A.U.)
Nor. TCR (A.U.)
**
ns
CMV(CueIL2)
CMV(noIL2) + FcIL2(Cue)
C
PCC IL2RA/TCR
PCC IL2RB/TCR
PCC IL2RG/TCR
****
D
PCC IL2RA/IL2RB
PCC IL2RA/IL2RG
PCC IL2RB/IL2RG
E
pSTAT5
pZAP70
Fc Cue IL-2
IST
CMV(Cue IL-2)
CMV(no IL-2) + FcCueIL-2
F
Nor. pSTAT5 (A.U.)
Nor. pZAP70 (A.U.)
G
PCC pSTAT5/pZAP70

Figure S6

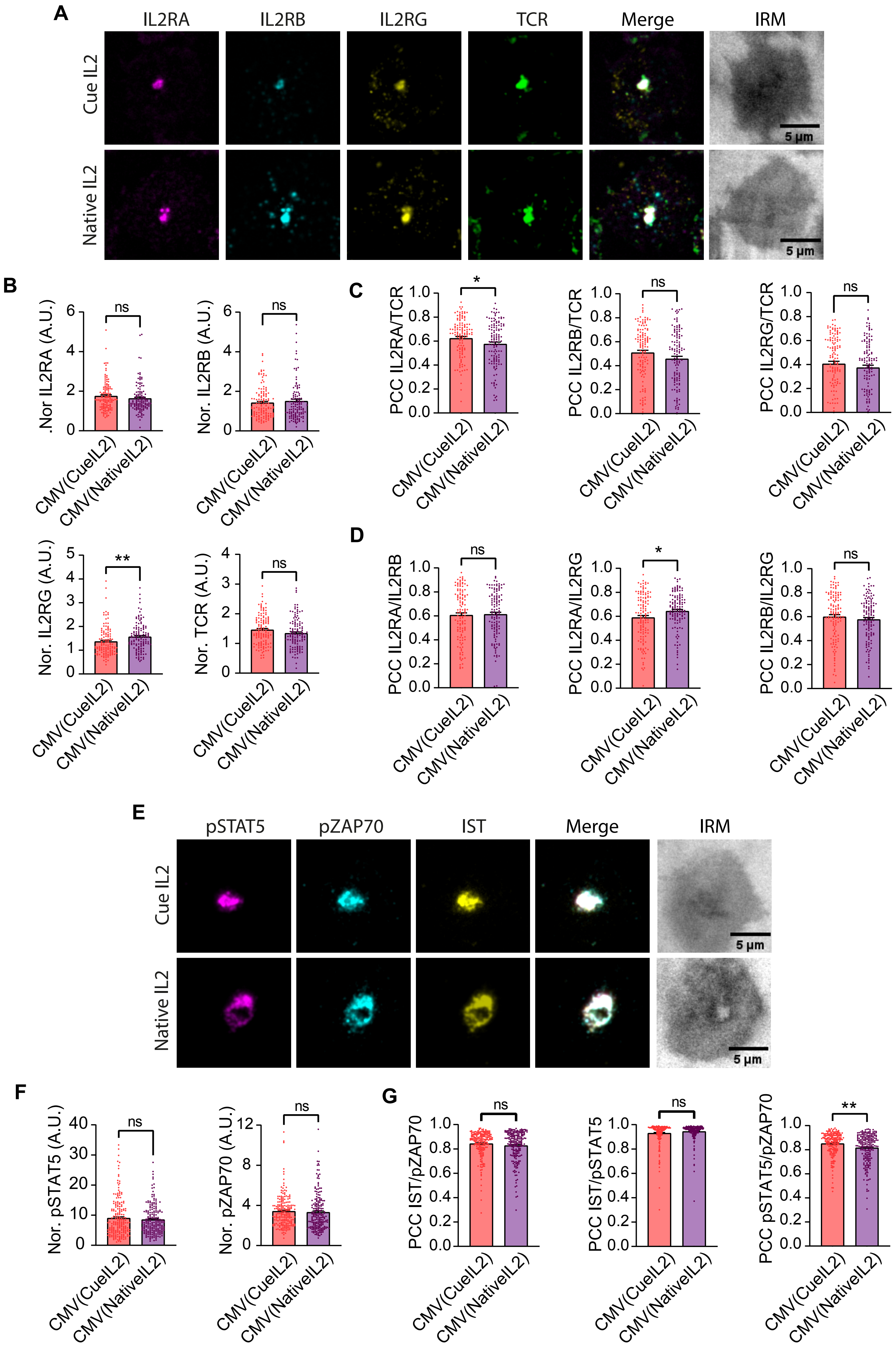
A
IL2RA
IL2RB
IL2RG
TCR
Merge
IRM
Cue IL2
Native IL2
5 µm
B
.Nor IL2RA (A.U.)
Nor. IL2RB (A.U.)
Nor. IL2RG (A.U.)
Nor. TCR (A.U.)
C
PCC IL2RA/TCR
PCC IL2RB/TCR
PCC IL2RG/TCR
D
PCC IL2RA/IL2RB
PCC IL2RA/IL2RG
PCC IL2RB/IL2RG
ns
*
**
CMV(CueIL2)
CMV(NativeIL2)
E
pSTAT5
pZAP70
IST
Merge
IRM
Cue IL2
Native IL2
F
Nor. pSTAT5 (A.U.)
Nor. pZAP70 (A.U.)
G
PCC IST/pZAP70
PCC IST/pSTAT5
PCC pSTAT5/pZAP70

Figure S7

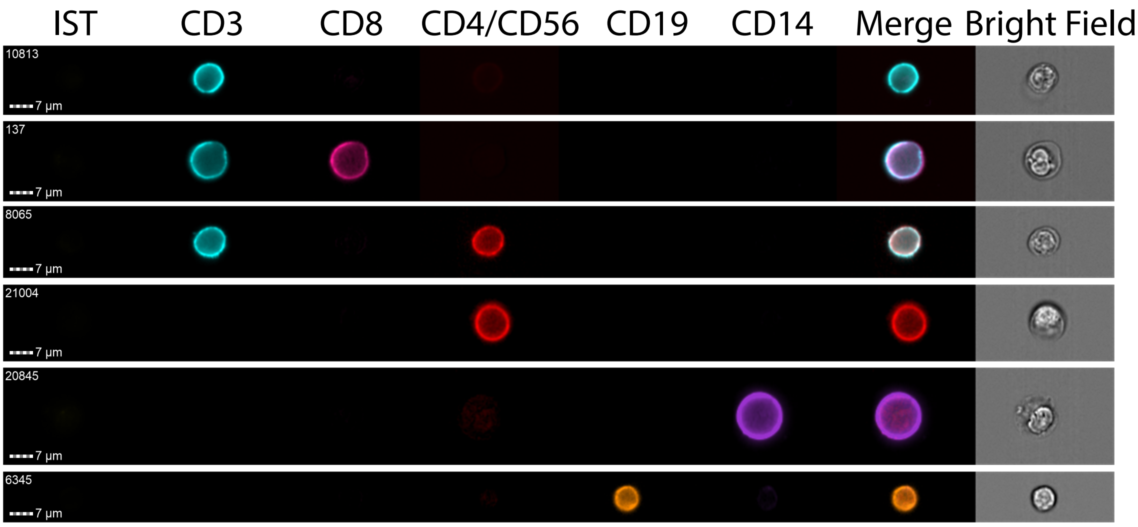
IST
CD3
CD8
CD4/CD56
CD19
CD14
Merge
Bright Field
10813
137
8065
21004
20845
6345
7 µm

Figure S8
